# *Shank3* loss heightens social engagement among sibling cagemates and antagonism toward strangers

**DOI:** 10.64898/2026.11.23.768613

**Authors:** Olivia Timmermans, Daniel Needs, Jenna McHenry

## Abstract

Autism spectrum disorder is a heterogeneous neurodevelopmental condition with strong genetic contributions and diverse social impairments. However, it remains unclear how autism-linked mutations alter social behavior across social contexts and relationships. Using SLEAP multi-animal pose tracking, we followed complete *Shank3* knockout (exon 4-22 deletion) mice with adult male and female sibling cagemates over four weeks and with strangers. Unexpectedly, knockouts initiated more social contact and allogrooming with familiar cagemates. Individual differences in social behavior and preferences for specific cagemates remained stable over time across genotypes. With strangers, knockout mice showed less nose-to-nose contact, more nose-to-base contact, and more chasing. Unlike wild-type mice, knockouts did not increase contact with strangers relative to siblings. These genotype differences were not detected in conventional three-chamber sociability and social novelty assays. These findings reveal that *Shank3* loss reshapes social behavior in context-specific ways, enhancing engagement with familiar partners while altering threat responses to social novelty.

## Introduction

Autism spectrum disorder (ASD) is a highly heritable and heterogeneous neurodevelopmental condition marked by atypical social communication, repetitive behaviors, and difficulties with emotional and homeostatic regulation^1,2^. Its clinical presentation varies widely across individuals, reflecting diverse genetic and environmental contributions. Among the genetic risk factors, mutations in *SHANK3,* a synaptic scaffolding gene associated with Phelan-McDermid syndrome (PMDS), represent a well-established monogenic cause of ASD^3,4^. Loss of SHANK3 disrupts synaptic organization and plasticity and has been associated with altered social behavior, anxiety-like phenotypes, and repetitive behaviors in both humans and animal models^5–7^. However, whether *Shank3* mutations produce generalized reductions in sociability or selectively alter specific components of social behavior across contexts remains poorly understood.

Historically, characterization of social behavior in *Shank3*-deficient mice has relied heavily on conventional assays, particularly the three-chamber sociability and social novelty tests^8,9^. Although these approaches have identified social phenotypes, they provide limited insight into the dynamic, reciprocal, and partner-specific features of naturalistic social interactions^10^. Moreover, conventional measures of social preference may obscure meaningful differences in how animals initiate, receive, and reciprocate social contact, or respond to familiar versus unfamiliar partners. Manual scoring of these interactions is labor-intensive and often limited to a coarse set of predefined behaviors, further constraining the resolution of social phenotyping.

To overcome these limitations, we employed Social LEAP Estimates Animal Poses (SLEAP), an open-source, deep-learning-based platform for markerless multi-animal pose tracking^11^. We developed a seven-point body-tracking model that maintains individual identities across visually indistinguishable mice, enabling automated, high-resolution quantification of social interactions. We applied this approach to a *Shank3* Δe4–22 mouse model, in which deletion of exons 4–22 disrupts all major SHANK3 isoforms^12^. Because Phelan-McDermid syndrome results from loss of one copy of SHANK3^4^, we compared heterozygous and knockout mice with their wild-type siblings. Using adult male and female mice, we examined social behavior across familiar sibling groups through repeated testing over four weeks, acute encounters with novel stranger conspecifics, and conventional behavioral assays, including anxiety-like assays and the three-chamber sociability and social novelty preference test. Together, this approach allowed us to determine whether *Shank3* loss broadly impairs sociability or instead produces context-dependent shifts in social engagement, partner-directed behaviors, and social interaction dynamics, and to compare how well naturalistic and conventional assays distinguish genotypes.

## Results

### *Shank3* loss enhances social familiarity dynamics

To quantify naturalistic social interactions, we established an automated machine-learning pipeline using SLEAP to track multiple mice with high spatial and temporal resolution. We developed a custom seven-point body skeleton enabling simultaneous markerless tracking of individual identity and full-body pose in groups of up to five mice (**Fig. 1a–c**; **Video S1**), with high model accuracy (**Fig. S1a–c**). Using head orientation and nose-centered contact windows, we extracted individual social encounters at 40 fps (**Fig. 1d**) and generated pairwise social contact matrices encompassing 147 behavioral metrics, including temporal and spatial features of social engagement (**Fig. 1e**; **Table 1**).

**Figure 1.**
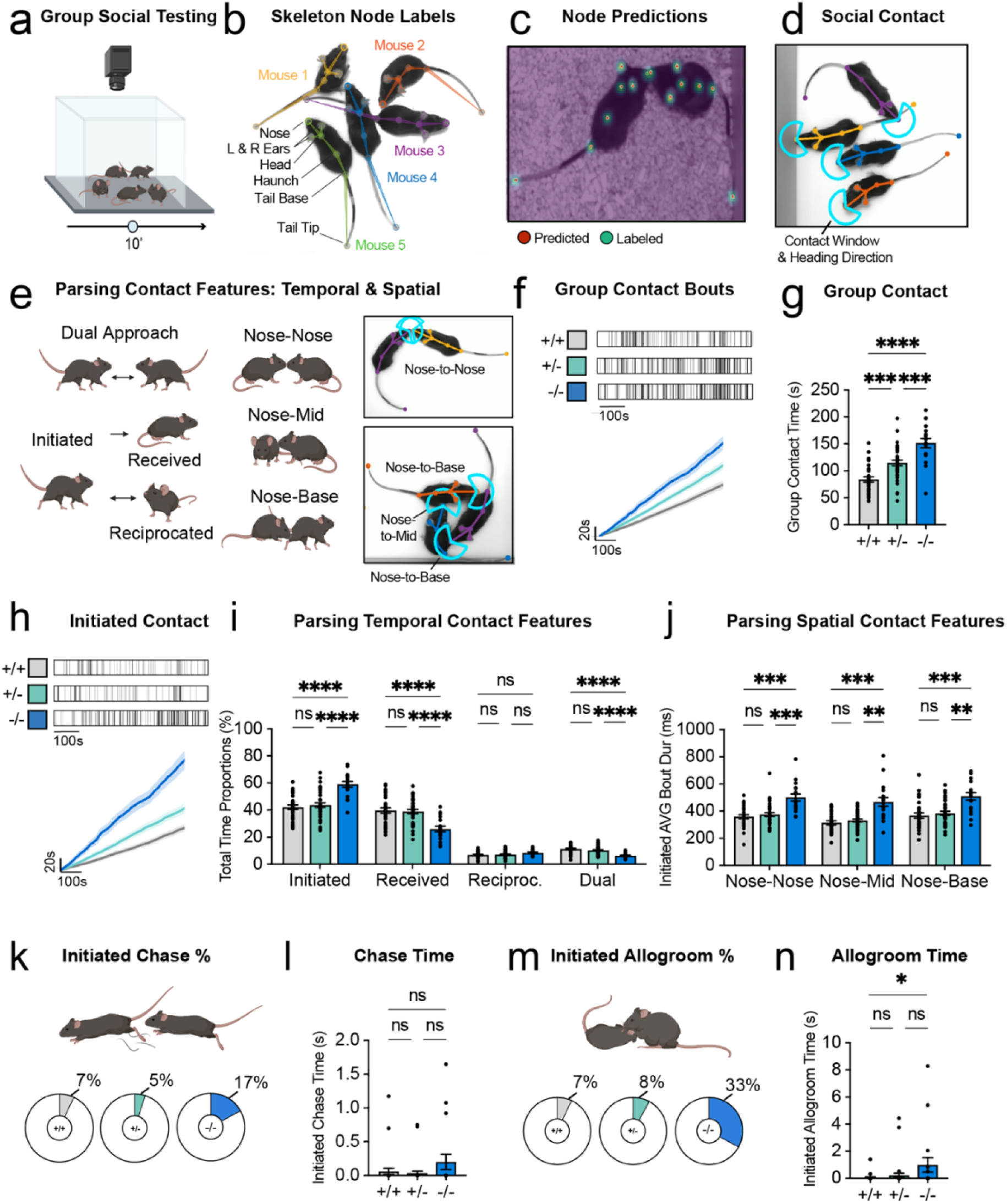
*Shank3* Δe4–22 knockout mice show elevated social engagement with familiar cagemate siblings. (A) Group social testing. Same-sex sibling groups housed together since weaning (2–5 mice per cage) were recorded from above during 10 min of free interaction in a 40 × 40 cm open-field arena. (B) Seven-point SLEAP skeleton (nose, left and right ears, head, haunch, tail base, tail tip), with individual identities maintained across five interacting mice. (C) Example frame showing model-predicted (red) and manually labeled (green) keypoints. (D) Social contact detection (blue). A 240° contact window (2 cm radius, ±120° about the heading direction) was centered on each mouse’s nose; contact was scored when this window intersected another mouse’s skeleton. (E) Contact bouts classified by temporal role (initiated, received, reciprocated, or dual approach) and by the body region contacted on the target mouse (nose-to-nose, nose-to-mid, or nose-to-base), with SLEAP examples. (F) Representative group-contact rasters (top; scale bar, 100 s) and cumulative group contact time (s) over the session (bottom). (G) Total group contact time. (H) Representative initiated-contact rasters (top; scale bar, 100 s) and cumulative initiated contact time (s) (bottom). (I) Proportion of total contact time that was initiated, received, reciprocated, or dual. (J) Mean duration of initiated contact bouts by spatial contact type. (K) Percentage of mice in each genotype that initiated chasing (WT 7%, 2 of 30; HET 5%, 2 of 38; KO 17%, 3 of 18). (L) Initiated chase time. (M) Percentage of mice in each genotype that initiated allogrooming (WT 7%, 2 of 30; HET 8%, 3 of 38; KO 33%, 6 of 18). (N) Initiated allogrooming time. WT, wild-type (Δe4–22^+/+^); HET, heterozygous (Δe4–22^+/−^); KO, knockout (Δe4–22^−/−^). Genotype colors: WT, gray; HET, teal; KO, blue. Data are represented as mean ± SEM with individual mice as dots; shading in cumulative plots shows ± SEM. n = 30 WT, 38 HET, and 18 KO mice from 27 cages (20 males and 10 females WT, 12 and 26 HET, 8 and 10 KO); males and females combined. One-way ANOVA with Tukey’s post hoc test (G, L, N); two-way repeated-measures ANOVA with Tukey’s post hoc test (I, J). Brackets: long bar, WT versus KO; short bars, WT versus HET (left) and HET versus KO (right). *p < 0.05, **p < 0.01, ***p < 0.001, ****p < 0.0001; ns, not significant. See also Figures S1 and S2.

**Table 1.** Automated SLEAP social contact and locomotor measures.

| Category | Measure (units) | Level | Definition |
|---|---|---|---|
| Locomotion | Average speed (cm/s) | Individual | Mean speed of the centroid across the session, from its displacement over 8-frame (200-ms) intervals. |
| Locomotion | Distance traveled (cm) | Individual | Speed integrated over the session (trapezoidal rule). |
| Spatial occupancy | Time in corner, at edge, and in center | Individual | The arena was divided into a 4 × 4 grid of equal 10 × 10-cm zones. Proportion of session time the centroid was in the 4 corner zones, the 8 non-corner edge zones, or the central 2 × 2 block (20 × 20 cm). |
| Contact detection | Contact | Directed pair | Frame on which the focal mouse’s contact window (240° sector, 2-cm radius, centered on the nose and aligned with the heading direction) intersected another mouse’s skeleton (ear–head, nose–head, head–haunch, and haunch–tail-base edges, 0.25 cm wide; tail excluded). |
| Contact group | Group | Individual | Mice linked directly or indirectly by contact in either direction; used to measure group size (time in group of k, see below). |
| Temporal role | Initiated contact | Directed pair | The focal mouse’s own contact brought the pair into contact (or it continued contacting after its partner stopped). Roles are assigned separately for each pair of mice. |
| Temporal role | Received contact | Directed pair | The partner initiated and the focal mouse is not contacting it on that frame. |
| Temporal role | Reciprocated contact | Directed pair | The partner initiated and the focal mouse is also contacting it, having joined on a later frame. |
| Temporal role | Dual contact | Directed pair | Both mice first contacted each other on the same frame. |
| Temporal role | Total contact (s) | Individual | Union of frames in which the focal mouse gave or received contact with any cagemate; equals the sum of initiated, received, reciprocated, and dual contact. |
| Spatial location | Nose-to-nose, nose-to-mid, nose-to-base contact | Directed pair | The partner’s nose, centroid, or tail base, respectively, was closest to the nose of the contacting mouse (assessed on the first frame of each period of unchanged contact). |
| Contact summary | Contact time (s) | Individual; directed pair | Cumulative time in each temporal role × spatial location combination, including totals. |
| Contact summary | Bout count | Individual; directed pair | Number of bouts; a bout is a run of consecutive frames with the same temporal role and spatial location. |
| Contact summary | Mean bout duration (ms) | Individual; directed pair | Summed bout durations divided by bout count; bout duration = number of frames × 25 ms. |
| Contact summary | Bout duration SEM (ms) | Individual; directed pair | Sample standard deviation (n − 1) of bout durations divided by the square root of bout count. |
| Cage-relative | Cage contribution | Individual | Contact time divided by the summed contact time of all mice in the cage. |
| Cage-relative | Cage-normalized contact | Individual | Contact time divided by the highest individual contact time in the cage. |
| Cage-relative | Min–max scaled contact | Individual | Contact time rescaled between the cage minimum (0) and maximum (1). |
| Pairwise | Normalized pairwise contact | Directed pair | Time in contact with one cagemate (given or received) divided by the focal mouse’s summed contact time with all cagemates. |
| Group state | Time in group of k (s) | Individual | Time during which the focal mouse’s contact group contained k mice; k = 1 indicates no contact. |

*Individual-level measures assign each mouse one temporal role at a time (priority initiated > dual > reciprocated > received), so simultaneous contacts are not double-counted. Pairwise measures are calculated for each pair from the focal mouse’s perspective and need not sum to individual totals. No gap bridging or minimum bout duration was applied. Equations are given in Supplementary Table S1.*

We first applied this approach to adult sex-and age-matched sibling groups housed together since weaning, containing wild-type (WT; Δe4–22⁺/⁺), heterozygous (HET; Δe4–22⁺/⁻), and knockout (KO; Δe4–22⁻/⁻) mice in their natural genotype distributions. Following habituation to a standard open-field arena, we examined spontaneous group interactions among cagemates at a single time point to establish baseline genotype differences. Unexpectedly, Δe4–22⁻/⁻ mice spent significantly more time in social contact with their cagemates than either heterozygous or wild-type siblings, with heterozygous mice intermediate (**Fig. 1f, g**). Parsing temporal contact features revealed that Δe4–22⁻/⁻ mice spent a greater proportion of time initiating contact, but less time receiving or engaging in simultaneous bidirectional (dual) contact, with no difference in reciprocated contact (**Fig. 1h, i**). Knockout mice also exhibited longer initiated contact bouts across spatial contact types, despite similar relative distributions of nose-to-nose, nose-to-mid, and nose-to-base interactions across genotypes (**Fig. 1j**; **S1d**).

We next examined more complex social behaviors. Although chasing did not differ across genotypes and no fighting was observed, Δe4–22⁻/⁻ mice initiated more prosocial allogrooming (**Fig. 1k–n**). Mice that chased moved about three times faster during chases than their session average in every genotype (Fig. **S1e**). The amount of chasing and allogrooming received did not differ by genotype (**Fig. S1f, g**). Outside social interactions, Δe4–22⁻/⁻ mice showed increased self-grooming and reduced locomotion, with no differences in center time relative to wild-type controls (**Fig. S1h–k**). Importantly, increased social contact and initiation persisted when comparisons were restricted to equal cage sizes or normalized for cage size, supporting the robustness of these effects despite natural differences in sibling group composition (**Fig. S1l–o**). Further, these effects were sustained throughout the 10-min session, with no differences in initiated contact between the first and second halves of the session (**Fig. S1p**).

We lastly compared output metrics between male and female mice. Sex differences were largely absent across social and movement metrics, except for higher reciprocated contact in heterozygous males than in heterozygous females, and higher allogrooming in knockout males than in knockout females (**Fig. S2a–k**). No sex differences were detected in self-grooming, movement, or speed within genotypes, though both heterozygous and knockout males spent more time in the center than females of their respective genotype, and center time was similar between sexes in wild-type mice (**Fig. S2l–o**). Together, these findings reveal robustly elevated social engagement in Shank3-deficient mice among familiar cagemate siblings, largely characterized by increased contact initiation, prolonged interactions, and prosocial allogrooming, despite reduced locomotor activity.

### *Shank3*-elevated sibling dynamics are persistent

To determine whether genotype-dependent social behaviors persist over time, we repeatedly recorded sibling groups across four consecutive days and one session per week for the following three weeks using the same arena and SLEAP pipeline (**Fig. 2a**). Importantly, Δe4–22⁻/⁻ mice consistently exhibited greater total and initiated social contact than heterozygous and wild-type siblings across all time points, both within individual cages and across the cohort (**Fig. 2b–e**; **S3a, b**). Heterozygous mice also had more total contact than wild-type mice (**Fig. 2c**). Consistent with our initial findings, Δe4– 22⁻/⁻ mice received less contact and initiated more allogrooming, while chasing remained unchanged (Fig. S3c–h). Knockout mice also exhibited increased self-grooming and reduced locomotion throughout testing (**Fig. S3i–l**).

**Figure 2.**
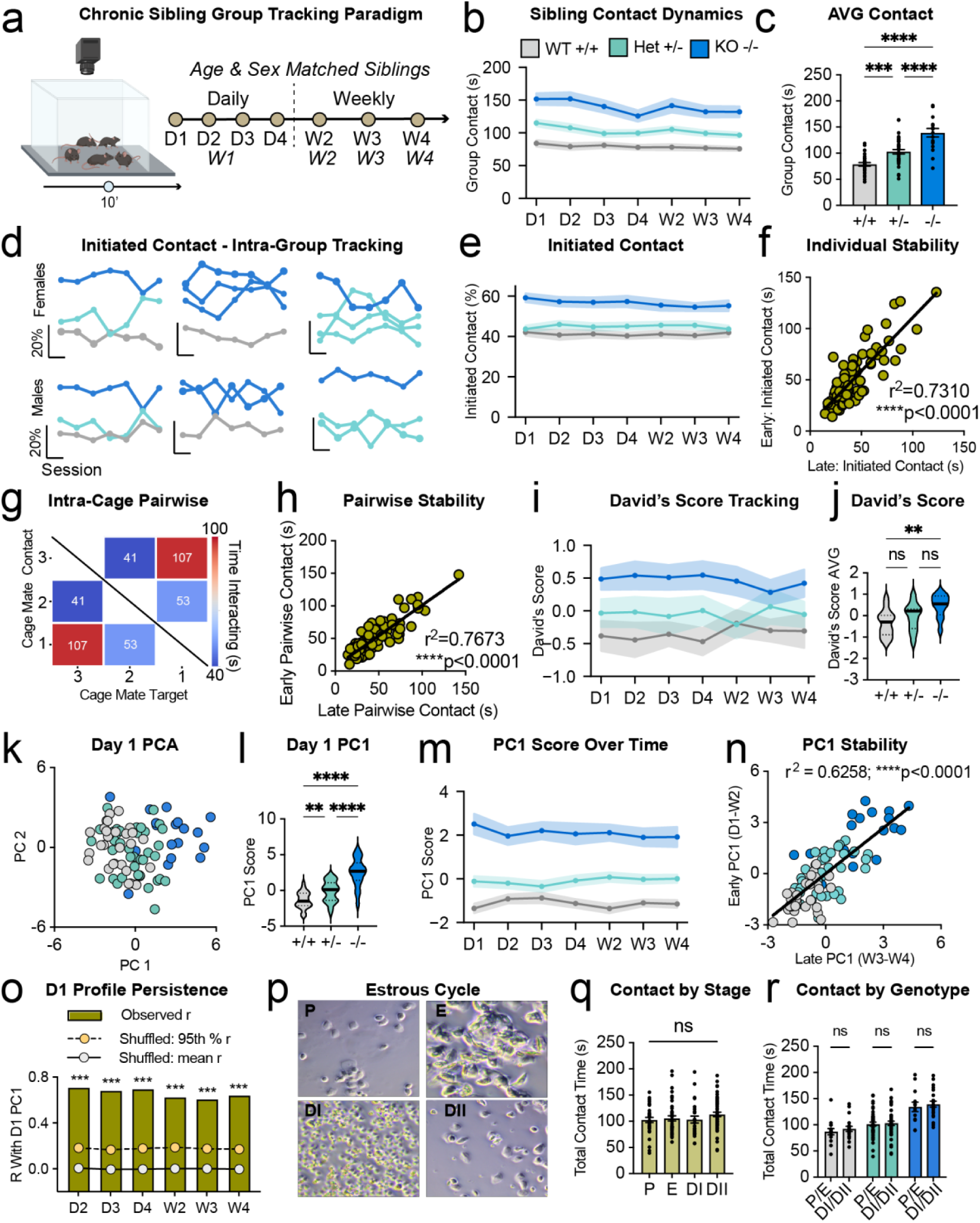
Elevated social engagement in knockout mice is stable over time and within individuals. (A) Chronic sibling-group tracking paradigm. Age-and sex-matched sibling groups were recorded for 10 min daily on four consecutive days (D1–D4; week 1) and then once weekly in weeks 2–4 (W2– W4), for seven sessions. (B) Group contact time across sessions. (C) Group contact time averaged across sessions. (D) Initiated contact of individual mice across sessions in three representative female (top) and male (bottom) cages; each line is one mouse, colored by genotype. Scale bars: 20% (vertical) and one session (horizontal). (E) Percentage of each mouse’s total contact time that it initiated, across sessions. (F) Individual stability: initiated contact time of each mouse in early versus late sessions (r² = 0.7310, p < 0.0001). (G) Intra-cage pairwise contact in a representative cage of three: total interaction time (s) for each pair of cagemates on a single day. (H) Pairwise stability: total contact time of each cagemate pair in early versus late sessions (r² = 0.7673, p < 0.0001). (I) David’s scores across sessions, calculated from directed initiated nose-to-base contact bouts (cages of three mice). (J) David’s score averaged across the seven sessions (higher values indicate higher rank). (K) Principal-component analysis (PCA) of 17 D1 sibling-group measures (Table S2), with each mouse colored by genotype; PC1 and PC2 explained 24% and 19% of the variance. (L) D1 PC1 scores (F(2,83) = 37.9, p < 0.0001). (M) PC1 scores at each session, with each later session projected onto the D1 principal components (genotype F(2,83) = 60.5, p < 0.0001; genotype × session F(12,498) = 1.20, p = 0.28). (N) Individual stability of PC1: early (mean of D1 and W2) versus late (mean of W3 and W4) PC1 scores (r² = 0.6258, p < 0.0001); dots are colored by genotype, and the line is fitted to all mice. (O) Persistence of D1 profiles: Pearson correlation (r) between each mouse’s D1 PC1 score and its PC1 score at each later session (bars; D2 0.71, D3 0.68, D4 0.69, W2 0.62, W3 0.61, W4 0.64), compared with the mean (white) and 95th percentile (yellow) of 1,000 per-measure shuffles of that session (mean ≈ 0; 95th percentile 0.17–0.19). (P) Representative vaginal cytology from females in proestrus (P), estrus (E), metestrus (diestrus I; DI), and diestrus (diestrus II; DII). (Q) Total contact time in females by estrous stage, pooled across genotypes. (R) Total contact time in females, with observations grouped as P/E or DI/DII within each genotype. Early and late values were the means of D1 and W2 and of W3 and W4, respectively. Lines and shading (or error bars in M) show mean ± SEM; bars show mean ± SEM with individual mice as dots; violin plots show the median (solid line) and quartiles (dotted lines). Lines in (F), (H), and (N) are linear regressions. n = 30 WT, 38 HET, and 18 KO mice (B–E, K–O); n = 86 mice, pooled genotypes (F, N regression); n = 105 cagemate pairs, pooled genotypes (H); n = 14 WT, 13 HET, and 12 KO mice in cages of three (I, J); n = 46 females with 33, 40, 26, and 58 sessions in P, E, DI, and DII, respectively, pooled genotypes (P, Q); 15 and 19 WT, 45 and 43 HET, and 13 and 22 KO observations in P/E and DI/DII, respectively (R). Two-way repeated-measures ANOVA (genotype × session) (B, E, I, M); one-way ANOVA with Tukey’s post hoc test (C, J, L, Q); simple linear regression (F, H, N); permutation test against 1,000 shuffles for each session, ***p = 0.001, the smallest attainable value (O); two-way ANOVA with Tukey’s post hoc test (R). Because each session was z-scored separately before projection, (M) tests genotype differences and genotype × session interaction, not overall change across sessions. Brackets: long bar, WT versus KO; short bars, WT versus HET (left) and HET versus KO (right). *p < 0.05, **p < 0.01, ***p < 0.001, ****p < 0.0001; ns, not significant. See also Figures S3 and S4.

Beyond genotype differences, individual and pairwise social relationships remained remarkably stable over time. Initiated contact was strongly correlated between early (W1 and W2) and late (W3 and W4) testing across the cohort (r² = 0.73; **Fig. 2f**) and within each genotype (r² = 0.24–0.65; **Fig. S3n–p**), and between every pair of sessions (r = 0.74–0.86; **Fig. S3m**). Similarly, each cage contained cagemate pairs with consistently high or low contact (**Fig. 2g**), and contact between specific pairs was highly correlated between early and late testing (r² = 0.77, n = 105 pairs; **Fig. 2h**), between every pair of sessions (r = 0.71–0.85; **Fig. S3q**), and within each genotype (r² = 0.69–0.73; **Fig. S3r–t**).

Collectively, these findings reveal persistent individual differences and partner-specific patterns of social engagement within sibling groups.

To assess whether social hierarchy contributed to these dynamics, we calculated David’s scores from directed nose-to-base pairwise contact, scoring each uninterrupted bout as a win for the initiating mouse, in cages of three mice (to avoid cage size biases in rank). Δe4–22⁻/⁻ mice had higher David’s scores than wild-type cagemates across sessions, with heterozygous mice intermediate and not different from either group, suggesting increased dominance-like or close investigatory follow behavior despite the absence of overt fighting (**Fig. 2i, j**).

To determine whether multidimensional social profiles distinguished genotypes and persisted across sessions, we applied principal-component analysis (PCA) to 17 Day 1 sibling-group measures selected without reference to genotype, excluding measures that varied with cage size or were redundant (**Table S2**). Genotypes separated along the first principal component (PC1; 24% of variance; **Fig. 2k**), which was weighted toward total and initiated contact, initiated bout duration, and the proportion of contact initiated. PC1 scores differed among all three genotypes, with heterozygous mice intermediate between wild-type and knockout mice (one-way ANOVA, F(2, 83) = 37.9, p = 2 × 10⁻¹²; **Fig. 2l**). When each later session was scored with the Day 1 PCA, the genotype difference persisted at every session (genotype F(2, 83) = 60.5, p = 6 × 10⁻¹⁷; genotype × session p = 0.28; **Fig. 2m**), and each mouse’s PC1 score was strongly correlated between early and late testing (r² = 0.63, p < 0.0001; **Fig. 2n**), including within each genotype (r² = 0.25–0.37, all p < 0.01). Each mouse’s Day 1 profile remained correlated with its profile at every later session (r = 0.61–0.71), far above the correlations obtained after shuffling each measure across mice (95th percentile, r ≈ 0.18; p = 0.001 for each session; **Fig. 2o**). Thus, an individual’s multidimensional social profile, and not only its initiated contact, was stable over four weeks.

Lastly, in female mice, we examined whether natural estrous cycling influenced social behavior during repeated testing by monitoring vaginal cytology in 46 females across 157 observations (**Fig. 2p**). Total contact did not differ across estrous stages, either pooled across genotypes or within each genotype (**Fig. 2q, r**). No differences were observed in initiated, nose-to-nose, or nose-to-base contact, or in average speed or center time (**Fig. S4a–j).** Together, these findings demonstrate that Shank3 loss produces persistent increases in social engagement and dominance-like behavior while preserving stable individual and partner-specific social relationships.

### *Shank3* loss disrupts encounters with novel partners

Next, we screened social interactions with an age-and sex-matched unfamiliar conspecific and a non-social novel object using the same SLEAP pipeline, providing a contrasting context to the established social dynamics among sibling cagemates (**Fig. 3a**; **Video S1, S2).** While total dyadic interaction time did not differ across genotypes, parsing temporal features revealed that Δe4–22⁻/⁻ mice initiated a greater proportion of social contact and spent less time receiving contact than either of the ot her genotypes (**Fig. 3b–f**). Analysis of spatial interaction patterns revealed that Δe4–22⁻/⁻ mice initiated more nose-to-base orientations, with longer average durations, and fewer nose-to-nose interactions than wild-type controls (**Fig. 3g–i**). Nose-to-nose interactions are often associated with prosocial behavior, whereas nose-to-base orientations may reflect antagonistic interactions, suggesting that Shank3 loss may reduce prosocial engagement and promote antagonistic behavior toward unfamiliar conspecifics. Indeed, genotype was a significant predictor of initiated chase (χ²(2) = 11.05, p = 0.004), with 89% of Δe4–22⁻/⁻ and 55% of Δe4–22⁺/⁻ mice displaying chase, compared with 40% of wild-type mice (**Fig. 3j)**. Furthermore, Δe4–22⁻/⁻ mice exhibited greater cumulative duration and frequency of initiated chasing toward the stimulus mouse, while also receiving less chasing from the stimulus mouse, compared with wild-type subjects (**Fig. 3k–m**; **Fig. S5c**). No genotype differences were detected in prosocial allogrooming (**Fig. S5a, b),** or in the duration of time spent interacting with a non-social novel object (**Fig. 3n**). Stereotypically elevated self-grooming was detected in Δe4–22⁻/⁻ mice during the novel object trial, but not during the social trial, together with more center time during the object trial and less locomotion (**Fig. S5d, e, h–j).**

**Figure 3.**
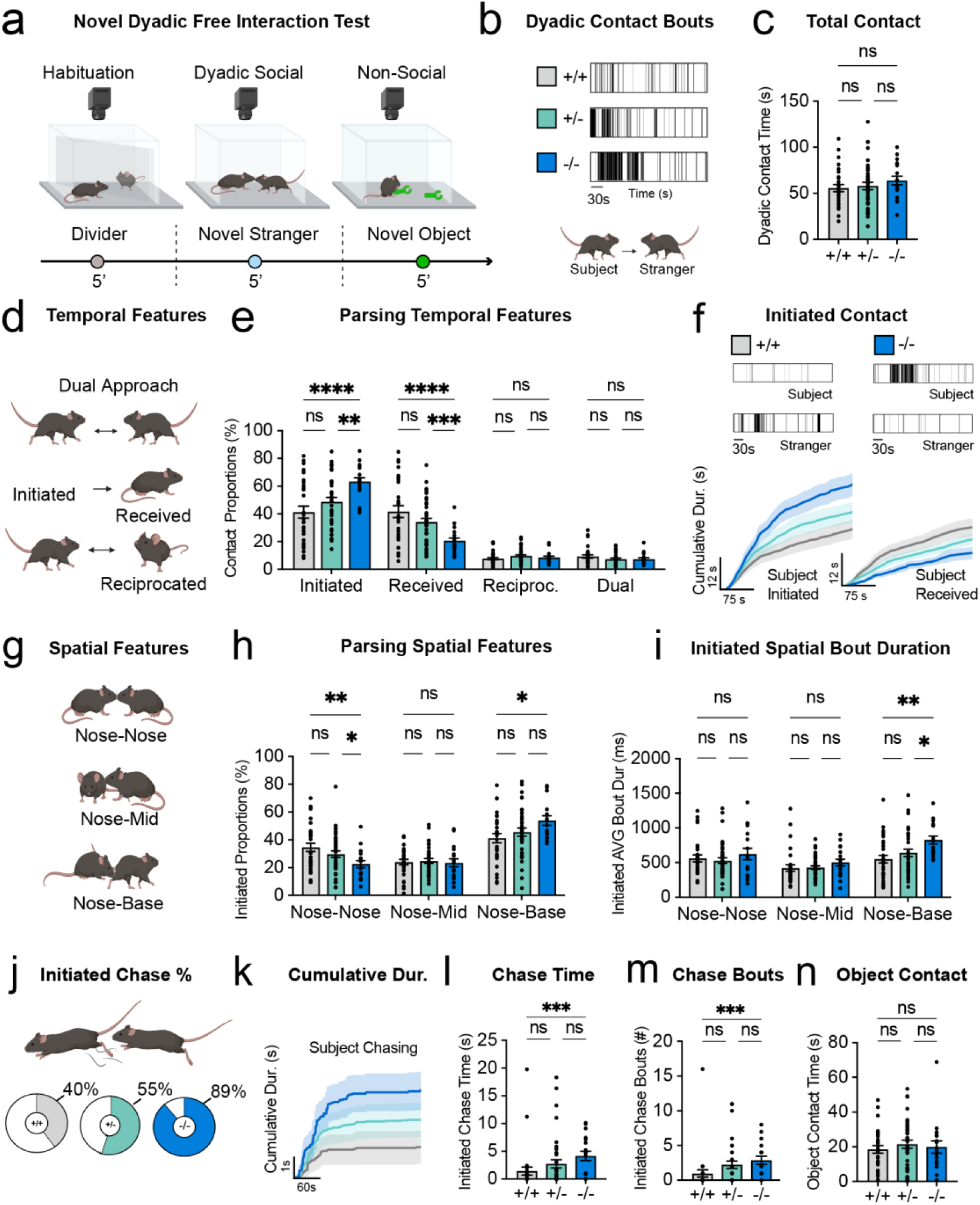
Knockout mice display altered behavior toward stranger mice. (A) Novel dyadic free-interaction test. After 5 min of habituation with a clear diagonal divider, the subject interacted freely for 5 min with an unfamiliar age-and sex-matched stranger. The stranger was then removed, and the subject explored two novel non-social objects (Nylabones) for 5 min. (B) Representative rasters of subject-to-stranger total contact (scale bar, 30 s). (C) Total dyadic contact time. (D) Temporal contact categories. (E) Proportion of total contact time that was initiated, received, reciprocated, or dual. (F) Representative rasters of contact initiated by a KO and a WT subject and by their strangers (top; scale bars, 30 s), and cumulative contact initiated (bottom left) and received (bottom right) by subjects over the session. (G) Spatial contact categories. (H) Proportion of initiated contact by spatial type. (I) Mean duration of initiated bouts by spatial type. (J) Percentage of subjects that initiated chasing the stranger (WT 40%, 12 of 30; HET 55%, 21 of 38; KO 89%, 16 of 18; χ²(2) = 11.05, p = 0.004). (K) Cumulative initiated chase duration over the session. (L) Initiated chase time. (M) Number of initiated chase bouts. (N) Time spent contacting the novel objects in the non-social trial. Data are represented as mean ± SEM with individual mice as dots; shading shows ± SEM. n = 30 WT, 38 HET, and 18 KO subjects (the same mice as in Figure 1). One-way ANOVA with Tukey’s post hoc test (C); two-way repeated-measures ANOVA with Fisher’s LSD test (E, H, I); chi-square test (J); Kruskal–Wallis test with Dunn’s multiple comparisons (L–N). Brackets: long bar, WT versus KO; short bars, WT versus HET (left) and HET versus KO (right). *p < 0.05, **p < 0.01, ***p < 0.001, ****p < 0.0001; ns, not significant. See also Figures S5 and S6.

To assess whether the observed phenotypes were sustained across the session as the subject became familiar with the stranger, we time-binned the data. Total, initiated, and chase contact and locomotion were higher in the first half of the session than in the second half in most genotypes, whereas received contact declined only in wild-type and heterozygous mice and object contact did not change (**Fig. S5k–r**). Together, these findings indicate that disrupted social novelty engagement observed in Δe4–22⁻/⁻ mice is generally restricted to the onset of the session, as the animal navigates social uncertainty or a potential threat response. Unlike sibling contexts, sex differences were detected across several temporal and spatial features of novel social encounters. Wild-type and heterozygous males initiated a greater proportion of contact and more nose-to-base contact than females of the same genotype, whereas females made proportionally more nose-to-nose contact, and heterozygous females received and reciprocated more contact (**Fig. S6b–f**). Wild-type females also reciprocated more contact than wild-type males, and heterozygous males allogroomed strangers more than heterozygous females (**Fig. S6d, h**). Knockout males spent more time in the arena center than knockout females, whereas heterozygous females spent more time in the center than heterozygous males, and no sex differences were detected in speed or distance moved (**Fig. S6m–o).**

### *Shank3* loss alters discrimination between novel and familiar partners

To assess social discrimination between familiar and novel social partners, we compared each subject’s dyadic behavior toward a stranger with its behavior toward its familiar siblings during the first 5 min of the Day 1 sibling session, an equivalent testing duration (**Fig. 4a**). Initiated contact did not differ between the first and second 5 min of that session (**Fig. S1p**). Contact times were not normalized for the number of available partners, because contact time did not vary with cage size.

**Figure 4.**
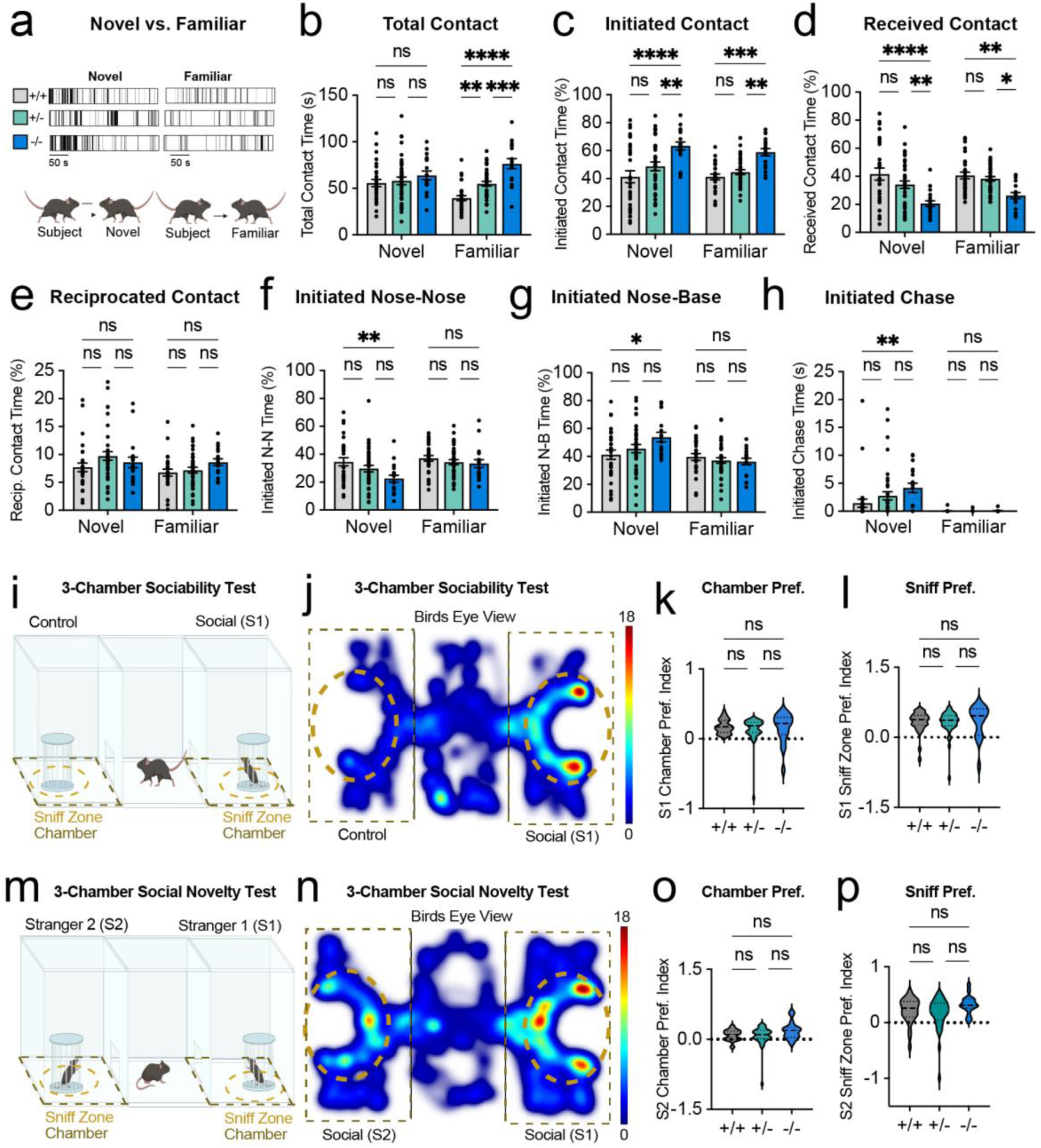
*Shank3* loss alters responses to novel versus familiar partners. (A) Each subject’s contact with a novel stranger (Novel; 5-min novel dyadic test) was compared with its contact with familiar cagemate siblings (Familiar; first 5 min of the D1 sibling-group session). Representative rasters for each genotype (scale bars, 50 s). (B–H) Genotypes compared within each partner condition: total contact time (B), percentage of contact time initiated (C), percentage of contact time received (D), percentage of contact time reciprocated (E), percentage of initiated contact that was nose-to-nose (F) or nose-to-base (G), and initiated chase time (H). (I) Three-chamber sociability test, with a novel stranger (S1) under a wire enclosure in one side chamber and an empty enclosure (control) in the other; dashed lines mark the chambers and sniff zones. (J) Occupancy heatmap during the sociability phase (representative mouse). (K and L) Sociability preference index for chamber (K) and sniff-zone (L) time, (T_S1_ − T_Baseline_)/(T_S1_ + T_Baseline_), where T_Baseline_ is time in the same chamber or sniff zone during habituation. (M) Social novelty test, with a second novel stranger (S2) placed in the previously empty enclosure, opposite the now-familiar S1. (N) Occupancy heatmap during the social novelty phase (representative mouse). (O and P) Social novelty preference index for chamber (O) and sniff-zone (P) time, (T_S2_ − T_Baseline_)/(T_S2_ + T_Baseline_). Contact times are raw values, not normalized for the number of available partners; the familiar condition is a group of cagemates and the novel condition a single partner (see STAR Methods). Data are represented as mean ± SEM with individual mice as dots; violin plots show the median (solid line) and quartiles (dotted lines); the horizontal dotted line at 0 marks no preference relative to habituation. n = 30 WT, 38 HET, and 18 KO mice (B–H); n = 29 WT, 38 HET, and 18 KO mice (K, L, O, P). Two-way repeated-measures ANOVA (genotype × partner) with Tukey’s post hoc test comparing genotypes within each partner condition (B–H); one-way ANOVA with Tukey’s post hoc test (K, L, O, P). Brackets: long bar, WT versus KO; short bars, WT versus HET (left) and HET versus KO (right). *p < 0.05, **p < 0.01, ***p < 0.001, ****p < 0.0001; ns, not significant. Novel-versus-familiar comparisons within each genotype are shown in Figure S7. See also Figures S7 and S8.

Wild-type mice spent more time in contact with novel strangers than with familiar siblings, whereas heterozygous and knockout mice did not (**Fig. S7a**). This difference arose from the familiar context: Δe4–22⁻/⁻ mice spent more time in contact with siblings than both other genotypes, whereas contact with strangers did not differ across genotypes (**Fig. 4b**). In both novel and familiar contexts, Δe4–22⁻/⁻ mice initiated more contact and received a smaller proportion of contact than the other genotypes (**Fig. 4c, d**), and proportions of initiated and received contact did not differ between contexts within any genotype (**Fig. S7b, c**). Reciprocated contact did not differ across genotypes (**Fig. 4e**), although heterozygous mice reciprocated more contact with novel partners (**Fig. S7d**). Importantly, toward strangers, but not siblings, Δe4–22⁻/⁻ mice showed less nose-to-nose and more nose-to-base contact than wild-type mice (**Fig. 4f, g**), and they showed less nose-to-nose and more nose-to-base contact with strangers than with siblings, as did heterozygous mice for nose-to-base contact (**Fig. S7e, f**).

Consistent with this pattern, Δe4–22⁻/⁻ mice chased strangers more than wild-type mice did (**Fig. 4h**), and both Δe4–22⁻/⁻ and Δe4–22⁺/⁻ mice chased novel strangers but not familiar siblings, a difference not detected in wild-type controls (**Fig. S7g**), suggesting heightened aggressive or threat-related responses specifically toward unfamiliar social partners as a result of Shank3 loss. Increased allogrooming of novel compared with familiar partners was detected only in heterozygous mice (**Fig. S7h**).

To contextualize these findings using a conventional behavioral assay, we employed the three-chamber sociability and social novelty test (**Fig. 4i, j**). During the sociability phase, chamber and sniff-zone preferences, calculated relative to each mouse’s habituation baseline, did not differ across genotypes (**Fig. 4k, l**), with all groups preferring social over non-social zones (**Fig. S7k, l).** Similarly, during the social novelty phase, no overall genotype differences were detected (**Fig. 4o, p**), and all groups preferred the novel social stranger (S2) over the familiar stranger (S1) **(Fig. S7m, n**).

Consistent with other assays, Δe4–22⁻/⁻ mice exhibited reduced locomotion compared with wild-type controls during the habituation and social novelty phases (**Fig. S7o–q).**

Given the time-dependent effects observed in the dyadic stranger assay, we further examined behavior across the first and second halves of each testing phase. During sociability testing, no genotype differences were detected in the first half of the session, but in the second half, Δe4–22⁻/⁻ mice spent more time with the social stimulus than both other genotypes (**Fig. S8a).** Within genotypes, both wild-type and heterozygous mice spent more time with the novel stranger in the first versus second half of the session, with no differences in knockout mice (**Fig. S8b).** No differences were detected between or within genotypes with respect to the non-social object (**Fig. S8c, d).** No genotype differences were detected between the first and second halves of the social novelty phase, though all genotypes spent more time with the S2 stranger in the first half (**Fig. S8e, f).** Lastly, both heterozygous and knockout mice spent more time with the S1 stranger in the second half of the session than in the first, an effect not observed in wild-type mice (**Fig. S8g, h).** No sex differences were observed in social preference during either sociability or social novelty testing (**Fig. S8i–l).**

### *Shank3* loss alters anxiety-like behavior and locomotor activity

To assess anxiety-like behavior and locomotion, we employed the open field test (OFT), elevated plus maze (EPM), and elevated zero maze (EZM) (**Fig. 5a, f, k**). In the OFT, Δe4–22⁻/⁻ mice exhibited reduced locomotion and center entries, but no differences in center time, including during the first five minutes of testing (**Fig. 5c–e**; **S9a–h)**. In contrast, both Δe4–22⁻/⁻ and Δe4–22⁺/⁻ mice exhibited increased open-arm exploration and reduced anxiety-like behavior in the EPM (**Fig. 5h–j**; **S9i, j**).

**Figure 5.**
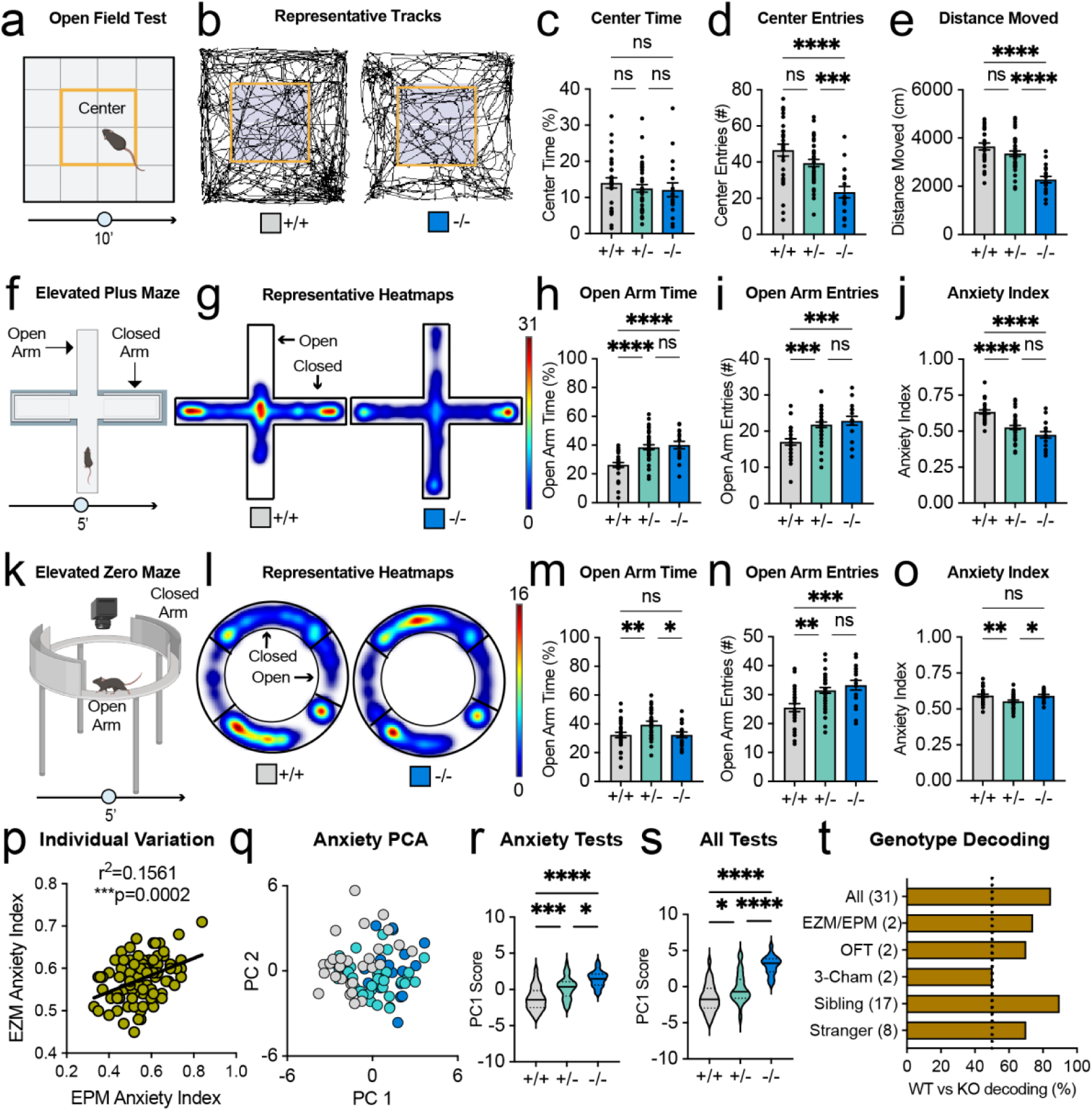
*Shank3* loss reduces locomotion and anxiety-like behavior, and combined behavioral profiles distinguish genotypes. (A) Open-field test (OFT; 10 min), with the center zone outlined. (B) Representative WT and KO tracks. (C) Percentage of time in the center. (D) Center entries. (E) Distance moved. (F) Elevated plus maze (EPM; 5 min). (G) Representative WT and KO occupancy heatmaps. (H) Percentage of time in the open arms. (I) Open-arm entries. (J) Anxiety index, 1 − [(open-arm time/total time + open-arm entries/total entries)/2]; higher values indicate more anxiety-like behavior. (K) Elevated zero maze (EZM; 5 min). (L) Representative WT and KO occupancy heatmaps. (M) Percentage of time in the open quadrants. (N) Open-quadrant entries. (O) Anxiety index, calculated as in (J) from open-quadrant time and entries. (P) Correlation between EPM and EZM anxiety indices, pooled across genotypes (r² = 0.1561, p = 0.0002). (Q) PCA of 10 anxiety-like and locomotor measures from the OFT, EPM, and EZM (Table S2), with each mouse colored by genotype; PC1 and PC2 explained 31% and 26% of the variance. (R) PC1 scores from (Q) (F(2,82) = 18.8, p < 0.0001). (S) PC1 scores from a PCA of 31 measures from five assays combined (novel dyadic test, D1 sibling group, three-chamber test, OFT, and elevated mazes; Table S2), with assays unweighted; PC1 explained 20% of the variance (F(2,82) = 31.2, p < 0.0001). (T) Genotype decoding: cross-validated balanced accuracy of logistic-regression classifiers distinguishing WT from KO mice using measures from each assay alone or all assays combined (number of measures in parentheses): all assays (31), 85%; EZM and EPM (EZM/EPM, 2), 74%; OFT (2), 70%; three-chamber (2), 50%; D1 sibling group (17), 90%; novel stranger (8), 70%. The dotted line marks chance (50%). Data are represented as mean ± SEM with individual mice as dots; violin plots show the median (solid line) and quartiles (dotted lines); the line in (P) is a linear regression. n = 29 WT, 38 HET, and 18 KO mice (A–S) and 29 WT and 18 KO mice (T) (mouse 1404 excluded, euthanized before completing all assays). One-way ANOVA with Tukey’s post hoc test (C–E, H–J, M–O, R, S); Pearson correlation (P); leave-one-cage-out cross-validation, with the all-assay model tested against 200 permutations of genotype labels (p = 0.005) (T). Brackets: long bar, WT versus KO; short bars, WT versus HET (left) and HET versus KO (right). *p < 0.05, **p < 0.01, ***p < 0.001, ****p < 0.0001; ns, not significant. See also Figure S9.

Effects in the EZM were less consistent: Δe4–22⁺/⁻ mice spent more time in the open quadrants and had a lower anxiety index than both other genotypes, whereas Δe4–22⁻/⁻ mice made more open-quadrant entries but did not differ from wild-type mice in open-quadrant time or anxiety index (**Fig. 5m–o**; **S9k, l)**. Individual differences in anxiety-like behavior were consistent across assays, as evidenced by a significant positive correlation between EPM and EZM anxiety indices when data were pooled across genotypes (**Fig. 5p**). This relationship was also present within wild-type and heterozygous mice, with a similar but nonsignificant trend in knockout mice (**Fig. S9y–aa**). To further characterize individual differences, we applied PCA to 10 non-redundant anxiety-like and locomotor measures (**Table S2**). PC1, on which higher scores reflected more open-arm exploration and less open-field locomotion and center entry, separated the genotypes (**Fig. 5q**), and PC1 scores differed among all three genotypes, with heterozygous mice intermediate (F(2, 82) = 18.8, p = 2 × 10⁻⁷; **Fig. 5r**). We next combined 31 measures from five assays (novel dyadic encounters, Day 1 sibling interactions, the three-chamber test, the open field, and the elevated mazes; Table S2). PC1 of this combined analysis was weighted toward initiated contact with strangers and siblings and toward nose-to-base rather than nose-to-nose contact with strangers, and separated the genotypes more strongly (F(2, 82) = 31.2, p = 9 × 10⁻¹¹; **Fig. 5s**). To ask which assays carried this information, we trained classifiers to distinguish wild-type from knockout mice using the measures from each assay and tested them on cages withheld from training. Day 1 sibling-group measures identified genotype with 90% accuracy, the elevated mazes, open field, and dyadic stranger test with 70–74% accuracy, and the three-chamber preference measures performed at chance (50%; **Fig. 5t**). Combining all 31 measures (85%) did not improve on the sibling-group measures alone. Thus, home-cage social behavior with familiar siblings was the most sensitive indicator of Shank3 loss, whereas the conventional three-chamber test was the least.

Sex-dependent differences were detected in subsets of anxiety and locomotor measures that varied by assay **(Fig. S9m–x).** Specifically, knockout females spent less time in and made fewer entries into the OFT center than knockout males over the full session, and spent less time in the center during the first 5 min (**Fig. S9m, n, q**), with no sex differences in movement **(Fig. S9o, p).** In the EPM, knockout males spent more time in the open arms than knockout females (**Fig. S9s**), and knockout females had a higher anxiety index than knockout males (**Fig. S9u**), whereas in the EZM, wild-type females spent more time in and made more entries into the open quadrants and had a lower anxiety index than wild-type males (**Fig. S9v–x).**

## Discussion

The primary goal of the present study was to assess more nuanced and distinct aspects of social behavior across different social contexts and relationships, as humans also experience a diverse range of social contexts that could be differentially affected in autism spectrum disorders. We also examined selective preferences for specific cagemates in group-level interactions among siblings of both sexes. Specifically, we used multi-animal pose tracking to characterize social behavior in *Shank3* Δe4–22 mice across familiar and novel social contexts, alongside conventional sociability assays. Importantly, we found that *Shank3* loss did not produce a generalized reduction in social interactions, but rather context-dependent alterations in specific social behaviors. Unexpectedly, Δe4–22 knockout mice initiated more social contact and exhibited elevated initiated social contact behaviors among familiar sibling cagemates compared with wild-type mice. Furthermore, irrespective of genotype, we identified stable individual differences in social behavior and selective, persistent preferences for specific cagemates. In contrast, when encountering novel strangers, knockout mice exhibited increased threat-like behaviors, highlighting the context-dependent nature of their social phenotype. Despite reduced locomotor activity across most assays, knockout mice maintained robust social engagement. Notably, these behavioral differences were not detected using conventional three-chamber sociability and social novelty assays. Together, these findings demonstrate that *Shank3* loss alters specific components of social behavior in a context-dependent manner, highlighting the importance of examining social interactions and neurobiology across different partners and social environments.

### Heightened Sibling Engagement and Group Cagemate Dynamics

While prior studies of the Δe4–22 mouse line used in the present study are limited, many other *Shank3* mouse lines have been screened for social behavior, although most have relied on conventional assays, such as the three-chamber test to measure sociability and social novelty and the tube test to measure dominance. To our knowledge, no other studies to date have assessed *Shank3* sibling intra-cagemate dynamics using natural mixed genotype compositions. In a different study, Live Mouse Tracker, a depth-sensing camera system with RFID tagging and machine-learning identification, was used to assess group dynamics in adult female ProSAP2/Shank3Δ11 mice (exon 11 deletion)^13^. In contrast to our study, experimenters deliberately assembled mixed-genotype groups, comprising six groups of four mice (two wild-type and two *Shank3* knockout), for at least 3 weeks leading up to the experiment and they placed littermates in different groups when necessary. Mice were tracked continuously for three days, with most analyses limited to the first 23 h. *Shank3* knockout mice showed few differences from wild-type mice. Specifically, they spent less time moving alone and less time at the end of three-mouse "train" configurations, but other social events were not significantly affected. In pairs of *Shank3*Δ11 knockout females housed together for at least 3 weeks, total contact time was typical, but nose-to-nose and side-to-side contacts lasted longer, and ultrasonic vocalization patterns were altered^14^. In another study, Live Mouse Tracker was used to measure social behavior during ten-minute trials in 2-month-old Shank3B mice (exon 13–16 deletion) of both sexes, reared in same-genotype groups of 2-4 mice^15^. Contrary to our findings, knockout mice showed less nose contact with cagemates than wild-type mice. However, in a separate reciprocal interaction experiment, four weeks of social isolation increased nose-contact time in knockout mice, consistent with our observation that *Shank3* social phenotypes depend on social context. Similarly, when *Shank3* knockout mice were introduced as newcomers into groups of wild-type mice, homozygous knockouts engaged more in social interactions despite reduced locomotion, whereas heterozygous mice showed reduced interaction^16^. Differences across these studies may partially reflect social housing and group composition, as groups were either assembled by experimenters from non-littermates^13^ or housed by genotype^15^, whereas our mixed-genotype sibling groups were housed together since weaning.

Differences may also arise from the distinct Shank3 mutations and genetic backgrounds used across studies.

### Dominance Behavior and Allogrooming

Several studies have examined the relationship between *Shank3* loss and dominance behavior. In contrast to our findings, *Shank3*Δ9 knockout males showed a lower probability of winning against wild-type littermates in the conventional tube test, and reduced performance in a separate cooperation task, whereas *Shank2*Δ6–7 knockout males showed a higher winning probability and a nonsignificant trend toward improved cooperation^17^. Notably, that study tested only males, and tube test opponents were non-cagemate littermates. Specifically, mice were housed with same-genotype cagemates and tested against wild-type littermates from other cages, whereas our measures in both sexes were taken spontaneously among mixed-genotype sibling cagemates. In the original characterization of the Δe4– 22 line, dominance was assessed in same-genotype triads of unfamiliar adults by ranking each mouse by the number and direction of agonistic behaviors within the triad across a 6-day span, combining data from males and females^12^. Agonistic behaviors included climb-grooming along the back of a cagemate, chasing, mounting, biting, and fighting. Wild-type groups established stable hierarchies by day 1 that remained stable throughout the 6-day period, whereas 75% of knockout triads were unstable, either failing to maintain dominance across days or lacking an identifiable dominant animal.

In contrast to these studies, in our mixed-genotype sibling groups housed together since weaning, we observed no overt fighting, even among males, across any of the daily or weekly recordings. Because chasing was also rare outside of stranger encounters, we measured pairwise nose-to-base following among cagemates, which revealed higher dominance-like scores in knockout mice than in wild-type mice. One limitation is that nose-to-base following, although used in some studies as an index of dominance, is ambiguous and may instead reflect closer sniffing, social investigation, or altered social chemosensory processing. Consistent with this possibility, a smaller proportion of vomeronasal sensory neurons responded to urine in group-housed Shank3B knockout males than in wild-types^15^, raising the possibility that increased nose-to-base contact reflects compensatory chemosensory sampling rather than dominance.

Along these lines, allogrooming carries mixed interpretations of social intent and can reflect affiliative, consoling touch^18,19^ or imposed, dominance-related grooming^20,21^. In a *Shank3* point-mutation line (R1117X) mutant mice caused facial hair loss in wild-type cagemates through allogrooming and mutant males won more tube-test encounters against unfamiliar wild-type males^22^. Our observation that Δe4–22⁻/⁻ mice allogroomed sibling cagemates more and showed higher dominance-like scores without overt aggression is consistent with these findings, suggesting a dominance-related component, although an affiliative component cannot be excluded and the two may not be mutually exclusive. In contrast, a pose-tracking study of male *Shank3B* mice tested in pairs showed that allogrooming was less common in pairs containing knockouts^23^, indicating that increased allogrooming is not consistent across *Shank3* models and testing conditions.

In the social novelty test in the present study, chasing was more frequent than among siblings, and it was asymmetric. For instance, knockout mice were more likely to chase the stranger, chased it for longer and more often, and received less chasing from it than wild-type subjects, together with more nose-to-base contact, mostly early in the session. Because each subject met a single stranger once and were in a dyad setting, we did not derive hierarchy measures such as David’s scores for these dyads. However, this asymmetry suggests a dominance-like or antagonistic stance toward unfamiliar partners that contrasts with the absence of overt antagonism among siblings.

### Sociability and Novelty Preferences

Most studies of *Shank3* social behavior have assessed interactions with unfamiliar partners using free dyadic tests or the three-chamber arena, often within broader behavioral batteries focused on anxiety-like, repetitive, motor, and cognitive behaviors^12,24^. In the original characterization of the Δe4–22 line, free dyadic interactions with an unfamiliar C3H/HeJ male revealed that knockouts engaged in prolonged, non-reciprocated interactions compared to wild-type mice, whereas all genotypes spent similar amounts of time in bidirectional interactions with the stranger^12^. In contrast to our study, behavior was manually annotated by blinded observers, predating modern tracking and pose estimation methods. Furthermore, our stimulus mice were sex-and age-matched C57BL/6J mice, and their dyadic assay included only males that had been singly housed for more than 14 days before testing.

More commonly, the three-chamber assay has been used to screen the effects of *Shank3* loss on sociability and social novelty preference, with inconsistent results across lines and laboratories. In the *Shank3B* line, knockouts showed impaired sociability and social novelty preference^6^, and in the Shank3^E13^ line, heterozygous and knockout mice lacked social novelty preference and later showed social recognition deficits with a juvenile partner^25^. Using the three-chamber test, Gora et al.^15^ also found that *Shank3B* knockouts lacked social novelty preference, which was restored by chronic social isolation. However, in adult *Shank3B* mice of both sexes, three-chamber sociability was typical despite markedly reduced reciprocal interaction^26^, suggesting inconsistency even within the same line.

Because these effects are measured relative to wild-type baseline levels, which may vary across testing settings and stimulus strains, age, and contexts, such discrepancies are difficult to interpret.

More closely aligned with our findings, we detected no genotype differences in either phase of the three-chamber test in the complete knockout line, consistent with the intact sociability preference originally reported for this line^12^. Similarly, three-chamber social approach was not strongly affected in an independently generated complete Δ4–22 deletion line, although homozygous knockouts strongly avoided novel non-social objects^24^. Likewise, *Shank3*Δ9 mice showed normal sociability and social novelty recognition^17^. Together, these findings suggest that the three-chamber test, which measures preference for a caged partner, may be insensitive to the changes we observed during free interaction, where knockout mice could be distinguished from wild-type mice with 90% accuracy from sibling-group behavior but only at chance from three-chamber preference. A similar dependence on testing context was reported by Tang et al.^16^: free group interactions revealed behavioral differences in *Shank3* heterozygotes that were not detected by the three-chamber test, whereas homozygous knockouts showed reduced three-chamber sociability but increased durations of specific social interactions when introduced into wild-type groups.

### Context-Dependent Social Processing: Novelty, Familiarity, and Memory

In the present study, we tested the same individuals across a battery of social assays to compare how *Shank3* loss affects social behavior with familiar and unfamiliar partners, alongside conventional anxiety-like and locomotor assays. Although many *Shank3* studies have used comprehensive behavioral batteries, these have largely focused on anxiety-like, repetitive, motor, and cognitive behaviors^12,24^, with social behavior typically assessed only with unfamiliar partners. Our design instead allowed us to compare free interaction with sibling cagemates and with strangers within each animal.

Using this comparison, *Shank3* knockout mice failed to show the increased contact with strangers, relative to siblings, seen in wild-type mice. This suggests that *Shank3* loss does not globally diminish social interaction but may instead reorganize social engagement, biasing it toward familiar partners while eliciting threat-like responses toward strangers. Notably, social discrimination appeared intact, as all genotypes preferred the second stranger over the first in the three-chamber social novelty test. Thus, knockout mice could distinguish novel from familiar mice. In contrast, impaired social recognition has been reported in the Shank3^E13^ line^25^, social memory deficits in the *Shank3B* line^27^, and a reduced proportion of social memory neurons in ventral CA1 of *Shank3B* knockout mice^28^. Familiarity has also been shown to shape *Shank3* phenotypes at the level of context: in Shank3^ΔC/ΔC^ males, exposure to novel environments lacking familiar features produced lasting failure to engage with conspecifics and objects upon re-exposure, which was prevented by including familiar features during the initial exposure^29^. Our partner-specific findings also parallel clinical findings in autism showing that fusiform responses vary with face familiarity and identity, with preserved responses to personally familiar faces and children’s faces but reduced responses to unfamiliar adults^30,31^, suggesting that social differences may depend partly on partner characteristics, including familiarity, rather than being uniform across social contexts. Together, these findings support the possibility that *Shank3* loss alters the relative value of familiar and novel partners while sparing the ability to distinguish them.

In our study, responses to strangers were concentrated at the start of the encounter. In time-binned analyses of the free dyadic test, contact declined from the first to the second half of the session in all genotypes, and chasing was primarily confined to the first half in knockout and heterozygous mice. Furthermore, received contact declined in wild-type and heterozygous mice across the session, but not in knockouts. Because locomotion also declined across the free dyadic session, these changes likely reflect both general habituation and increasing familiarity with the stranger. However, contact with novel objects did not change across the session. Time-binned three-chamber data further supported a familiarity bias during the sociability phase, as wild-type and heterozygous mice showed reduced time with the stranger in the second half, whereas knockout mice did not. During the social novelty phase, knockout and heterozygous mice showed increased time with the now-familiar mouse in the second time bin. By contrast, contact with familiar siblings did not decline within the session and remained elevated in knockouts across four weeks. These early responses to strangers may reflect heightened reactivity to social uncertainty, consistent with escape behavior and avoidance of novel objects in a complete Shank3 deletion line^24^.

Outside of social contexts, *Shank3* loss reduced anxiety-like behavior in the elevated mazes.

Specifically, both heterozygous and knockout mice explored the open arms of the elevated plus maze more than wild-type mice, with weaker effects in the elevated zero maze. This differs from the increased anxiety-like behavior reported in *Shank3B⁻/⁻* mice^6^ and in earlier studies of Δe4–22 deletion models, in which knockout mice spent less time in the open-field center, were slower to enter an illuminated chamber^12^, and showed heightened anxiety-like behavior as adults in the elevated zero maze^32^. Consistent with those reports, our knockout mice made fewer open-field center entries and moved less, so the direction of the effect depended on the assay. Differences in the isoforms disrupted across *Shank3* lines^12,24^, as well as in age and testing conditions, may contribute. Notably, reduced anxiety-like behavior in the mazes suggests that the early antagonistic response of knockout mice to strangers is not simply generalized anxiety but is specific to social uncertainty. Finally, combining social and non-social measures separated the genotypes more strongly in our principal-component and classifier analyses, and sibling-group social behavior identified knockout mice most accurately, indicating that social behavior in a familiar setting carries information about *Shank3* loss that conventional anxiety and sociability assays capture only in part.

### Partner-Specific Social Preferences and Individual Trait Stability

To our knowledge, few, if any, studies have tracked the same individuals and groups of *Shank3* mice longitudinally. Here, we found that individual variation in social behavior was stable over time within individuals in all genotypes, with the highest correlation in knockouts, although this difference was not significant and sample sizes differed across genotypes. Extending this to multidimensional profiles using principal-component analysis, each mouse’s Day 1 PC1 position remained correlated with its position at every later session, genotype differences in PC1 persisted across sessions without a genotype × session interaction, and mice fell along a continuum rather than forming separate groups.

Similar trait-like individuality has been described in wild-type mice outside of autism models. Specifically, stable individual differences in exploratory patterns emerge among genetically identical mice sharing an environment^33^, and group-housed male mice show stable "identity domains" across their behavioral repertoire^34^, which parallel the persistence of our PCA profiles.

A key advantage of repeated multi-animal group tracking is the ability to measure intra-cage preferences for specific cagemates. We found stable selectivity: each cage contained pairs that interacted consistently more or less from early (Day 1 and week 2) to late (weeks 3 and 4) sessions, and this held within each genotype. Together, these findings indicate that mice exhibit consistent preferences for particular cagemates rather than only a general level of sociability, and that *Shank3* loss does not disrupt these preferences under our testing conditions. Such partner-specific consistency broadly parallels evidence that group-housed mice develop stable dominance relationships through repeated social interactions^35,36^. However, since sample sizes and genotype composition varied across cages, we could not rigorously test whether mice of particular genotypes preferred cagemates of particular genotypes.

### Female Estrous Cycle and Sex Comparisons

Since females were used in the present study, and the estrous cycle has been shown to influence behavior, we screened our mice for estrous phase to determine whether it explained the observed individual variation. Within and across genotypes, we observed no correlations between estrous stage and social metrics. Similarly, a study in wild-type C57BL/6J female mice used unsupervised machine learning (MoSeq) to show that stable individual differences, rather than estrous state, explained behavioral variation in the open field^37^.

Given the male bias reported in autism, we also compared males and females across assays. Sex differences were minimal among siblings, except that knockout males allogroomed cagemates more, which may relate to dominance hierarchies. Sex differences emerged mainly with strangers. Wild-type and heterozygous males initiated more contact than females, particularly nose-to-base contact, whereas females made proportionally more nose-to-nose contact. This pattern was absent in knockouts, suggesting that *Shank3* loss may disrupt typical sex-specific responses to unfamiliar partners. No sex differences were found in the three-chamber test, whereas knockout females showed more anxiety-like behavior than knockout males in the open field and elevated plus maze. Thus, sex differences depended on context, appearing mainly with unfamiliar partners and in anxiety assays rather than among familiar siblings. Many prior *Shank3* studies have used only males or combined sexes^12^. Our results, together with reports of no sex differences in three-chamber sociability in juvenile *Shank3*Δ4–22 mice^38^ and of sex-dependent motor and repetitive behavioral phenotypes in *Shank3*Δ11 mice^39^, highlight the value of testing both sexes across multiple social contexts.

### Conclusions, Limitations, and Future Directions

To conclude, our main finding is that *Shank3* loss robustly enhances initiated social contact among sibling cagemates. While agonistic behavior was largely absent among familiar siblings of both sexes, encounters with novel strangers reliably increased threat-and dominance-like behavior. Together, these findings add to a growing body of literature showing that *Shank3* loss does not produce a global social deficit but rather a distinct, context-specific alteration in how mice engage with home-cage siblings and novel strangers. Strengths of the present study include repeated acute and chronic tracking of sibling cagemates in natural genotype compositions, using SLEAP pose estimation with high temporal and spatial precision. Using this pipeline across social contexts in the same mice allowed direct comparisons between familiar and novel partners. Sibling-group measures classified knockouts with 90% accuracy, whereas three-chamber preference measures could not, similar to the normal three-chamber sociability but reduced reciprocal interaction reported in *Shank3B* mice^26^.

We also recognize several limitations of the present study. As with most assessments of social behavior, intent cannot be determined, and affiliative versus antagonistic behaviors are largely inferred from spatial pose, except for chasing, which is less ambiguous. Further, overhead video alone does not capture certain vertical postures, odor cues, or vocalizations. The latter may reveal additional social phenotypes, as altered vocalization patterns have been reported in *Shank3*Δ11 knockout pairs during free interaction^14^. Although mixed natural genotype composition is a strength, it also introduces variation in genotype composition and cage size that cannot be fully accounted for. However, both normalizing for cage size and restricting analyses to equal cage sizes revealed the same pattern of elevated contact in knockout mice. Another limitation is that stranger encounters were dyadic, whereas sibling groups provided more opportunities for contact. Future studies should use multiple novel partners, as well as repeated encounters with the same stranger over time as it becomes familiar.

Efforts to distinguish generalized anxiety from social anxiety across familiar and novel contexts would also help clarify these threat responses. Importantly, these findings open the door for preclinical studies investigating the neurobiology that governs distinct forms of social relationships in autism-related conditions. Ultimately, tracking social behavior across familiar and novel contexts can help reconcile inconsistent *Shank3* findings and refine how social differences in Phelan-McDermid syndrome and autism are described with respect to distinct subtypes of social behavior and social relationships.

### Resource availability

### Lead contact

Further information and resource requests should be directed to the lead contact, Jenna McHenry.

### Materials availability

This study did not generate new unique reagents.

### Data and code availability

Data and code reported in this paper will be deposited into a repository and be shared by the lead contact upon reasonable request following peer review.

## Acknowledgments

We thank Dr. Yong-Hui Jiang for providing the *Shank3* Δe4–22 founder mice and Dr. William Wetsel and the Duke Behavior Mouse Core for maintaining the line. We thank Erika Blaine for scoring a subset of the behavior videos, Carissa Hansen for assistance with three-chamber pilot testing, and Nicholas Lusk for writing the initial version of the SLEAP-based analysis code. This work was supported by a grant from the Simons Foundation [882291, JM] and funds from Duke University to the McHenry Lab.

## Author contributions

J.M. conceptualized and designed the study, obtained funding, and supervised the project. J.M. and O.T. performed the experiments and collected the data. J.M. labeled SLEAP frames and trained the pose model. D.N. developed the SLEAP-based analysis code, building on initial versions by N.L.; O.T. wrote figure-visualization code, compiled the data, ran the analysis pipeline, and verified input and output files. J.M. prepared the first drafts of the figures and paper. J.M. and O.T. analyzed and interpreted the data, assembled the figures, and revised the manuscript. All authors approved the final version.

## Declaration of generative AI and AI-assisted technologies in the writing process

During the preparation of this work, the authors used Claude (Anthropic) to edit text for clarity, to check consistency among figures, legends, code, and methods, and to assist in debugging and revising the analysis code. All code changes were reviewed and tested by the authors. After using this tool, the authors reviewed and edited the content as needed and take full responsibility for the content of the publication.

## Supplemental information

Document S1. Figures S1–S9.

Tables S1 and S2. Equations for SLEAP-derived measures (Table S1) and measures entered in each principal-component analysis and genotype-decoding model (Table S2), included at the end of this document.

**Supplementary Video 1. Representative Day 1 sibling-group session, related to** **Figure 1**. First 4 min of a Day 1 sibling-group session in the open-field arena, rendered with SLEAP pose tracking. Each mouse’s skeleton (nose, ears, head, haunch, tail base, and tail tip) is drawn in its own color (subject ID 920, blue = 921, WT; orange = HET; 922, yellow = KO). The cyan sector at each mouse’s nose is its contact window (240° sector, 2-cm radius), used to detect social contact. The label at the top left gives the mouse IDs, with an arrow from the mouse initiating contact to its partner (arrow color indicates spatial category: magenta = nose-base; yellow = nose-mid; white = nose-nose; a double-headed arrow indicates simultaneous bidirectional contact). The numbers at the top right give each mouse’s current zone in the 4 × 4 arena grid.

**Supplementary Video 2. Representative novel dyadic test, related to** **Figure 3**. Full 5-min trial of a Δe4–22⁻/⁻ subject mouse with an unfamiliar age-and sex-matched conspecific, rendered with SLEAP pose tracking. Each mouse’s skeleton (nose, ears, head, haunch, tail base, and tail tip) is drawn in its own color (Subject ID 1432, KO in blue, C57 stimulus 1324 stranger in orange). The cyan sector at each mouse’s nose is its contact window (240° sector, 2-cm radius), used to detect social contact. The label at the top left gives the mouse IDs, with an arrow from the mouse initiating contact to its partner (arrow color indicates spatial category: magenta = nose-base; yellow = nose-mid; white = nose-nose; a double-headed arrow indicates simultaneous bidirectional contact). The numbers at the top right give each mouse’s current zone in the 4 × 4 arena grid.

## STAR★METHODS

### KEY RESOURCES TABLE

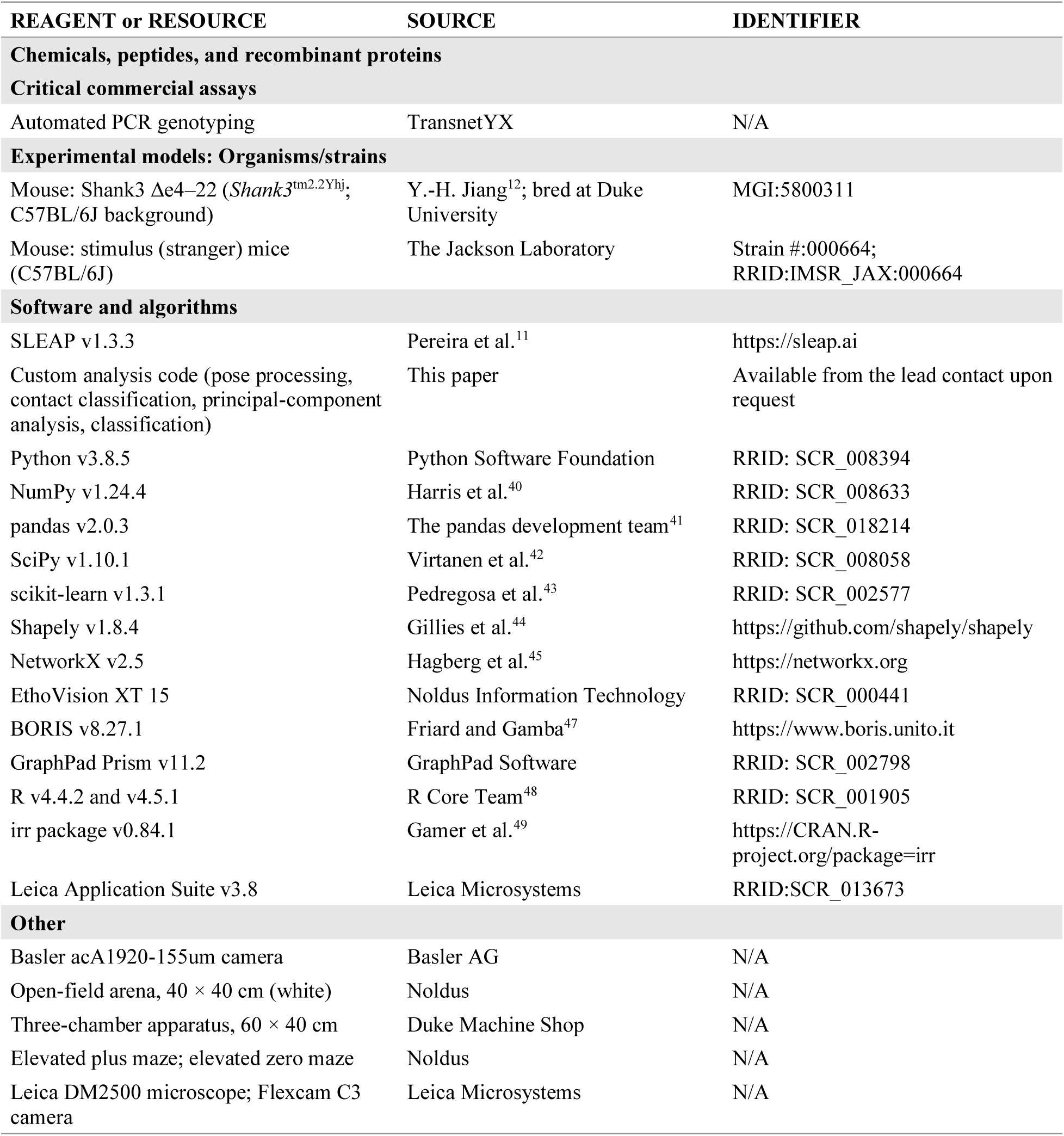

| REAGENT or RESOURCE | SOURCE | IDENTIFIER |
|---|---|---|
| **Chemicals, peptides, and recombinant proteins** |  |  |
| **Critical commercial assays** |  |  |
| Automated PCR genotyping | TransnetYX | N/A |
| **Experimental models: Organisms/strains** |  |  |
| Mouse: Shank3 Δe4–22 (*Shank3*<sup>tm2.2Yhj</sup>; C57BL/6J background) | Y.-H. Jiang<sup>12</sup>; bred at Duke University | MGI:5800311 |
| Mouse: stimulus (stranger) mice (C57BL/6J) | The Jackson Laboratory | Strain #:000664; RRID:IMSR_JAX:000664 |
| **Software and algorithms** |  |  |
| SLEAP v1.3.3 | Pereira et al.<sup>11</sup> | https://sleap.ai |
| Custom analysis code (pose processing, contact classification, principal-component analysis, classification) | This paper | Available from the lead contact upon request |
| Python v3.8.5 | Python Software Foundation | RRID: SCR_008394 |
| NumPy v1.24.4 | Harris et al.<sup>40</sup> | RRID: SCR_008633 |
| pandas v2.0.3 | The pandas development team<sup>41</sup> | RRID: SCR_018214 |
| SciPy v1.10.1 | Virtanen et al.<sup>42</sup> | RRID: SCR_008058 |
| scikit-learn v1.3.1 | Pedregosa et al.<sup>43</sup> | RRID: SCR_002577 |
| Shapely v1.8.4 | Gillies et al.<sup>44</sup> | https://github.com/shapely/shapely |
| NetworkX v2.5 | Hagberg et al.<sup>45</sup> | https://networkx.org |
| EthoVision XT 15 | Noldus Information Technology | RRID: SCR_000441 |
| BORIS v8.27.1 | Friard and Gamba<sup>47</sup> | https://www.boris.unito.it |
| GraphPad Prism v11.2 | GraphPad Software | RRID: SCR_002798 |
| R v4.4.2 and v4.5.1 | R Core Team<sup>48</sup> | RRID: SCR_001905 |
| irr package v0.84.1 | Gamer et al.<sup>49</sup> | https://CRAN.R-project.org/package=irr |
| Leica Application Suite v3.8 | Leica Microsystems | RRID:SCR_013673 |
| **Other** |  |  |
| Basler acA1920-155um camera | Basler AG | N/A |
| Open-field arena, 40 × 40 cm (white) | Noldus | N/A |
| Three-chamber apparatus, 60 × 40 cm | Duke Machine Shop | N/A |
| Elevated plus maze; elevated zero maze | Noldus | N/A |
| Leica DM2500 microscope; Flexcam C3 camera | Leica Microsystems | N/A |

### METHODS

#### Subjects

Shank3 Δe4–22 mice were bred in-house from founders originally provided by Dr. Yong-Hui Jiang^12^, and subsequently maintained at the Duke Behavior Mouse Core by Dr. William Wetsel. The line had been backcrossed for at least 12 generations onto a C57BL/6J background.

Genotypes (wild-type, heterozygous, and homozygous) were confirmed by automated PCR-based testing (TransnetYX). Both adult males and females were used throughout the study. Sibling-group tracking began at 9–27 weeks of age (median 14 weeks). Testing was replicated across 5 cohorts of mice. Cohort 5 was older at Day 1 (24–27 weeks) than cohorts 1–4 (9–17 weeks); however, initial analyses detected no age-related differences, so cohorts were combined. All genotypes were represented within each cohort. Sample sizes for each experiment are given in the figure legends. One cage of four mice was excluded from all analyses because one cagemate required euthanasia. Mice were housed with same-sex littermates from weaning (2–5 per cage), except when a sex-matched cagemate was unavailable. Adult mice were maintained on a reverse 12-h light/dark cycle (lights off at 06:00), with food and water available ad libitum. All procedures were approved by the Institutional Animal Care and Use Committee at Duke University and conducted in accordance with NIH guidelines for the care and use of laboratory animals.

#### General behavioral testing

Behaviorally naïve mice were acclimated to routine handling, body weights, and health checks in a colony room with a reverse light cycle for at least one week before assay habituation.

Behavioral testing occurred between 07:00 and 18:00 during the dark phase under dim red and infrared illumination, in a sound-attenuating testing environment. Arenas were cleaned with 70% ethanol between trials. Testing followed a consistent assay sequence across animals.

#### Chronic sibling-group tracking

Same-sex sibling groups were acclimated and tested in a 40 × 40-cm white open-field arena. Mice were habituated to handling, transport, tail marking, and the arena before testing, which consisted of ∼2 weeks of acclimation at increasing intervals, 2-3 days per week. Home-cage groups were transported in a blackout cart and acclimated to the testing room for at least 30 min prior to recording. Unique ear-punch identifiers and temporary tail marks supported video-based identity verification. Each cage of mice was recorded during free interactions in the field for 10 minutes per session on four consecutive days (D1–D4), followed by one session per week at weeks 2, 3, and 4 (W2–W4), for seven sessions in total; D1 served as the week-1 session.

Recordings used a Basler acA1920-155um camera at 40 frames/s and 1200 × 1200-pixel resolution. Sessions were conducted at approximately the same time of day during the dark cycle.

Unless otherwise stated, Day 1 (D1) analyses use the full 10-min D1 recording, and chronic summary values are the mean of the seven sessions. Early and late values for stability analyses were the mean of D1 and W2 and the mean of W3 and W4, respectively.

#### Estrous-cycle monitoring

For female mice, vaginal cytology was collected immediately after behavioral testing on sibling test days, following handling habituation as previously described^50^. A small volume (30 μL) of 0.9% sterile physiological saline was gently flushed at the vaginal opening without insertion, and the sample was examined by light microscopy to classify proestrus, estrus, metestrus (diestrus I), or diestrus (diestrus II) using cellular morphology (Leica Microsystems DM 2500, Flexcam C3, 10× magnification and phase contrast illumination). Images were captured using Leica Application Suite software (Version 3.8). Estrous stage was identified based on the morphology and density of characteristic cell types present in the vaginal secretion^50,51^, which reflect fluctuations in circulating gonadal steroids. Notably, the term "estrus" here refers to cytological staging, not to behavioral receptivity. Smears with insufficient cell yield or unclear classification were excluded from estrous-stage analyses.

Each female’s behavior in a given session was matched to the stage of the smear collected after that session. The final estrous dataset comprised of 157 observations from 46 females (proestrus 33, estrus 40, diestrus I 26, diestrus II 58).

#### Novel dyadic test

For novel dyadic interactions, each mouse was paired with an unfamiliar, age-and sex-matched conspecific of similar weight in a 40 × 40-cm arena. During a 5-min habituation period, a clear plexiglass divider was placed diagonally to prevent interaction. Then the divider was removed, and mice were free to interact for 5 minutes while overhead video was recorded for SLEAP analysis. Immediately afterward, the stimulus mouse was removed. The subject mouse remained in the arena and two novel non-social stimulus objects (2 green Nylabones) of similar size and color were placed in the arena (one centrally and one in a corner). The subject was free to interact with the objects for 5 minutes.

To compare contact with a familiar versus a novel partner (Fig. 4 and Fig. S7), each subject’s first 5 minutes of the D1 sibling-group session (familiar cagemates) was compared with its 5-min novel dyadic interaction (one unfamiliar partner). The same SLEAP pipeline and contact definitions were applied to both. Contact times are reported as raw values without normalization for the number of available partners: in the D1 group sessions, individual contact time did not differ among cages of 3, 4, and 5 mice, and dividing by the number of cagemates introduced a cage-size dependence that was absent from the raw data. Because the familiar condition involves several cagemates and the novel condition one partner, the comparison is between a familiar group and a novel dyad rather than between single partners. Genotypes were compared within each partner condition in Fig. 4b–h, and novel and familiar partners were compared within each genotype in Fig. S7a–h.

#### Three-chamber sociability and social novelty test

Mice were first habituated to a standard 60 x 40-cm arena empty three-chamber apparatus^8,9^ for 10 minutes. For sociability, an unfamiliar age-and sex-matched mouse (Stranger 1) was placed beneath a wire-mesh enclosure in one side chamber, opposite an empty enclosure, and the test mouse explored for 10 minutes. For social novelty, a second unfamiliar mouse (Stranger 2) was placed beneath the previously empty enclosure for another 10-min session. EthoVision XT 15 (Noldus^46^) quantified chamber occupancy, entries, and investigation of stimulus-enclosure sniff zones as shown in the figure diagrams. Sociability preference index = (TStranger1 − TEmpty)/ (TStranger1 + TEmpty); social novelty index = (TStranger2 − TStranger1)/ (TStranger2 + TStranger1). Indices were calculated for the stated chamber or sniff-zone measure. To rule out innate side biases, preference indices were also calculated from baseline as follows: Sociability preference index = (TStranger1 – TBaseline) / (TStranger1 + TBaseline); social novelty index = (TStranger2 – TBaseline) / (TStranger2 + TBaseline), where TBaseline is the time the same mouse spent in the corresponding chamber or sniff zone during habituation. Baseline-referenced indices are shown in Fig. 4k, l, o, p; raw chamber and sniff-zone times in Fig. S7i–n; and time-binned measures in Fig. S8.

#### Open-field and elevated maze tests

In the open-field test^52^, mice freely explored a 40 × 40-cm arena for 10 minutes. EthoVision XT 15 was used to quantify locomotion and center versus periphery occupancy. In the elevated plus maze^53^, mice were placed in the center facing an open arm and allowed to explore for 5 minutes; the apparatus included two open and two closed arms (30 × 5 cm), with 15-cm closed-arm walls, elevated 50 cm. In the elevated zero maze^54^, mice explored for 5 minutes on a circular track (60-cm outer diameter, 5-cm width), divided into alternating open and closed quadrants with 15-cm walls and elevated 50 cm. Time in open areas, entries, and distance traveled were quantified as appropriate using EthoVision XT 15. Open-area occupancy was used as an anxiety-like behavioral measure; closed-arm or closed-quadrant time and entries are shown in Fig. S9i–l.

Open-field measures were analyzed for the full 10 min and for the first 5 min. An anxiety index was calculated for each maze^55^ as 1 − [(open time/total time + open entries/total entries)/2], so that higher values indicate more anxiety-like behavior; the open-field index for the first 5 min (Fig. S9h) was calculated in the same way from center time and center entries.

#### SLEAP pose estimation

A custom bottom-up SLEAP^11^ model (v1.3.3) used a seven-keypoint skeleton (nose, bilateral ears, head, haunch, tail base, and tail tip) with a U-Net backbone (maximum stride 64, output stride 4, 32 base filters, filter rate 1.5, one stack) to track multiple interacting mice. The bottom-up multi-instance model predicted part confidence maps (σ = 1.5 pixels, output stride 4) and part affinity fields for six skeleton edges (σ = 30 pixels, output stride 8) from full-resolution grayscale frames (1200 × 1200 pixels; input scaling 1.0). Training used random rotation (±180°) as the only augmentation, the Adam optimizer (initial learning rate 1 × 10⁻⁴, halved after 5 epochs without improvement, minimum 1 × 10⁻⁸), a batch size of 4, and up to 200 epochs, with early stopping after 10 epochs without improvement in validation loss. Training frames were obtained from top-down videos recorded at 40 frames/s, including groups of interacting mice in arenas with and without bedding. A single annotator labeled 61,675 animal instances across 15,692 frames, which were split at the frame level into training (14,123 frames; 55,487 instances) and validation sets (1,569 frames, 10%; 6,188 instances). Frames were selected iteratively to represent difficult poses, including rearing, close contact, chasing, fighting, and allogrooming.

Occluded anatomical landmarks were annotated at their inferred positions to support learning of partially occluded poses.

#### SLEAP model evaluation

Performance was evaluated against manually annotated validation frames using object keypoint similarity (OKS), mean average precision (mAP), visible-keypoint precision and recall, and keypoint localization error. On the validation set, the final model achieved OKS mAP = 0.908, visible-keypoint precision = 0.9998, and visible-keypoint recall = 0.986. Median and mean keypoint localization errors were 1.12 and 2.34 pixels, and the 75th and 95th percentile errors were 2.00 and 6.06 pixels, respectively, in 1200 × 1200-pixel frames. Using an approximate spatial calibration of 25.24 pixels/cm derived from comparable recordings (D1) in the 40 × 40-cm arena, these correspond to approximately 0.044, 0.093, 0.079, and 0.240 cm, respectively.

#### Pose inference, identity tracking, and quality control

Videos were processed in batches with SLEAP command-line inference (sleap-track, v1.3.3) using the model described above and the simple tracker, with the target instance count set to the number of mice in each video (2–5). In each frame, detections were culled to that number (pre-cull IoU threshold = 2.0, so that culling was by count only) and linked across frames by bounding-box intersection-over-union (IoU) similarity with Hungarian matching over a 20-frame track window. Identity assignments and skeleton positions were manually inspected, and observed identity swaps and misplaced keypoints were corrected before export. Proofread tracks were analyzed for the first 10 min of each recording (or the full recording, if shorter). Frames in which a mouse or keypoint was missing were filled by linear interpolation of each coordinate, with no maximum gap length; missing values at the start or end of a recording were set to the nearest observed value. To estimate how much data were interpolated, missing keypoints were quantified in the analyzed windows of the D1 sibling-group sessions (27 videos) and novel dyadic tests (86 videos), excluding the tail tip, which was not used for contact detection. Missing keypoints accounted for 0.39% and 0.25% of keypoint-frames, respectively. Gaps longer than 10 frames (0.25 s) were rare (0.18% and 0.10% of keypoint-frames; median length 0.4 s); the longest were 6.5 s and 4.7 s, both at the tail base.

#### Automated social contact detection

Pose coordinates were exported from SLEAP and processed with custom Python scripts (Python v3.8.5; available from the lead contact upon request). The analysis scripts were debugged and revised with assistance from Claude (Anthropic), and all changes were reviewed and tested by the authors. Pixel coordinates were converted to centimeters using a scale derived from the four arena corners (40 × 40 cm), which were marked on a video frame of each recording. Each mouse’s centroid was defined as the midpoint between its head and haunch landmarks, and its heading as the direction of the sum of the vectors from the head landmark to the left and right ears. For each mouse, a contact window was defined as a 240° circular sector with a 2-cm radius, centered at the nose and oriented along the heading (±120°). Social contact was registered on each frame in which this window intersected another mouse’s skeleton, represented by the ear– head, nose–head, head–haunch, and haunch–tail-base edges, each 0.25 cm wide; the tail (tail base to tail tip) was excluded. Frame-level contact was stored in a directed mouse-by-mouse-by-frame matrix (40 frames/s; 25-ms resolution). No gap bridging, filtering, or minimum bout duration was applied.

#### Temporal roles and spatial locations of contact

Each contact was assigned a temporal role separately for each pair of mice, so that other cagemates did not change a pair’s roles. Contact was scored as initiated when the focal mouse’s own contact window brought the pair into contact; received when the focal mouse was approached and was not contacting its partner; reciprocated when the approached mouse also contacted its partner; and dual when both mice first contacted each other on the same frame.

Each contact was also assigned a spatial location (nose-to-nose, nose-to-mid, or nose-to-base) according to whether the partner’s nose, centroid, or tail base was closest to the nose of the contacting mouse. A bout was defined as a run of consecutive frames with the same temporal role and spatial location, and its duration as the number of frames × 25 ms. When contact became one-sided again (one mouse stopped contacting while the other continued), the mouse still contacting was scored as initiating a new contact and its partner as receiving it. Contact already in progress on the first frame of a recording was classified by the same rules. Contact groups (mice linked directly or indirectly by contact) were used only to measure group size. For each role × location combination, we derived cumulative contact time, bout count, mean bout duration and its SEM, and cage-relative contact measures.

At the individual level, a mouse in contact with several cagemates at once was assigned one role per frame, in the priority initiated > dual > reciprocated > received; when several cagemates shared that role, the spatial location for the first of them in track order was used. For each pair, spatial location was assessed on the first frame of each period in which the pair’s directed contact was unchanged, from the Euclidean distances between the contacting mouse’s nose and the partner’s nose, centroid, and tail base (for received contacts, the partner’s nose and the focal mouse’s landmarks). Bouts were defined separately for each temporal role × spatial location combination, so a continuous contact that changed role or location was counted as more than one bout. Bout duration SEM was calculated as the sample standard deviation (n − 1) divided by the square root of the bout count.

#### Individual and pairwise contact measures

Contact was summarized at the individual and pairwise levels. At the individual level, each mouse was assigned a single temporal role at any given time, so simultaneous contacts with several cagemates were not double counted; total contact was the union of all frames in which the mouse gave or received contact and equaled the sum of its four temporal roles. At the pairwise level, contact between the focal mouse and each cagemate (given or received) was summarized separately, so a mouse could participate in several dyadic contacts at once, and pairwise durations need not sum to individual total contact time. Centroid trajectories were used to calculate speed, distance traveled, and the proportion of time spent in the corner, edge, and center zones of a 4 × 4 grid. All measures are defined in Table 1; equations are given in Table S1.

Cage-relative measures divided each mouse’s contact time by the cage total (cage contribution) or the cage maximum (cage-normalized contact), or rescaled it between the cage minimum and maximum (min–max scaling). Normalized pairwise contact divided the time a focal mouse spent in contact with one cagemate (given or received) by its summed contact time with all cagemates. Time in a group of k mice was the time during which the focal mouse’s contact group contained k mice; k = 1 indicates no contact with any cagemate. In Fig. S1n,o, contact time was divided by the number of available partners (cage size − 1).

#### Locomotion and zone occupancy

Speed was calculated from the displacement of the centroid over 8-frame (200-ms) intervals, V(t) = ‖c(t) − c(t − 8)‖ · f / 8, where c is centroid position in centimeters (pixel coordinates divided by the x and y pixel-to-centimeter scales) and f = 40 frames/s; the first 8 frames took the first full-interval value. Average speed was the session mean of V(t), and distance traveled was V(t) integrated over the session (trapezoidal rule, Δt = 25 ms).

Zone occupancy used a 4 × 4 grid of equal 10 × 10-cm zones scaled to the four arena corners, comprising four corner zones, eight non-corner edge zones, and a 20 × 20-cm center zone formed by the four inner squares, and is expressed as the proportion of session time in which the centroid was in each zone type.

#### Manual annotation of complex behaviors

Behaviors not reliably captured by pose-derived contact measures were annotated from video in BORIS^47^ (v8.27.1). For social behaviors, initiator and recipient identities were recorded. Chasing was defined as rapid pursuit of another mouse that was actively fleeing. Mounting required forepaw clasping and thrusting; no mounting events were observed. Allogrooming was defined as licking or mouth contact directed toward another mouse’s trunk, shoulder, or head, distinguished from agonistic interactions by the absence of recipient flinching or avoidance. Self-grooming included licking or paw-directed grooming of the animal’s own body. Fighting was defined as attacks involving attempted biting or pinning; no fighting events were observed.

Chase speed was the mean centroid speed (calculated as described under Locomotion and zone occupancy) across the frames of each mouse’s annotated initiated chase bouts and was compared with that mouse’s session-average speed (Fig. S1e).

#### Validation of automated contact detection

To evaluate automated social-contact detection, a rater blinded to the automated output annotated contact bouts and their temporal and spatial features in BORIS. Automated and manually scored metrics were compared for three social-novelty trials (six mice total), including cumulative contact time, relative temporal and spatial contact proportions, and representative bout ethograms. Comparing manual and automated annotations, we assessed frame-by-frame agreement of raw contact labels with Cohen’s kappa and agreement of total contact time per trial with two-way intraclass correlation coefficients (ICCs; single-measures, absolute agreement and consistency), both calculated in R (v4.5.1) with the irr package (v0.84.1). Automated and manual contact detection agreed with a mean κ of 0.978 across the three trials (0.976, 0.974, and 0.984), and total contact time gave ICC (agreement) = 0.985 and ICC (consistency) = 0.999.

#### Longitudinal stability and dominance-like hierarchy

Individual stability was assessed for initiated contact time and pairwise stability for total contact time of each cagemate pair, by simple linear regression of early (mean of D1 and W2) on late (mean of W3 and W4) values, as plotted in Fig. 2f,h. Pearson correlations between every pair of the seven sessions summarized longitudinal relationships (Fig. S3m, q). Pairwise stability used 105 cagemate pairs (total contact in either direction; duplicate pairs removed); in the genotype-specific plots (Fig. S3r–t), each pair was included under the genotype of each of its members, so mixed-genotype pairs appear in more than one panel; because pairs from the same cage are not independent, pairwise regressions are descriptive.

Dominance-like relationships were estimated from directed nose-to-base contact. Each uninterrupted run of frames in which one mouse initiated nose-to-base contact with a cagemate was scored as one interaction won by the initiating mouse. For each session, interactions were summed into a win matrix for each cage, and David’s scores^56,57^ were calculated from dyadic dominance indices corrected for chance (Dij). David’s score analyses (Fig. 2i, j) were restricted to cages of three mice (39 mice: 14 wild-type, 13 heterozygous, 12 homozygous) so that the score range did not depend on group size, and each mouse’s David’s score was averaged across the seven sessions (Fig. 2j).

#### Principal-component analysis of behavioral profiles

Behavioral profiles were summarized by principal-component analysis (PCA) in Python (scikit-learn^43^; code available from the lead contact upon request). Three analyses were run: (i) the D1 sibling-group SLEAP measures (Fig. 2k–o), (ii) anxiety-like and locomotor measures from the open field, elevated plus maze, and elevated zero maze (Fig. 5q, r), and (iii) measures from all assays combined, including the novel dyadic test, D1 sibling group, three-chamber test, open field, and elevated mazes (Fig. 5s). The measures entered in each analysis are listed in Table S2. Measures were selected before analysis by rules that did not use genotype. For the SLEAP group measures, we excluded (1) measures calculated from cagemates’ values (cage contribution, cage-normalized and min–max scaled contact) and time in a group of k, whose possible values depend on cage size; (2) measures that were zero for most mice; (3) measures that differed among cages of 2–5 mice (cage size as a factor in a genotype-adjusted ANOVA, p < 0.05), because cage size differed among cages and would otherwise drive the principal components; and (4) one of each pair of near-duplicate measures (|r| > 0.9). This left 17 D1 measures covering the amount of contact, temporal role, spatial location, bout duration, and arena use. For the anxiety analysis, near-duplicate EthoVision measures (|r| > 0.9) were removed, leaving 10. For the all-assay analysis, 31 measures were drawn from five assays (novel dyadic 8, D1 sibling group 17, three-chamber 2, open field 2, elevated mazes 2). Mouse 1404 was excluded from analyses ii and iii (euthanized before completing all assays) (85 mice: 29 wild-type, 38 heterozygous, 18 homozygous); analysis i included 86 mice (30, 38, and 18).

Each measure was z-scored across mice, and PCA was fitted to the z-scores; measures were not weighted by assay. Because the sign of a principal component is arbitrary, the first and second components (PC1 and PC2) were oriented so that the knockout mean was positive. PC1 explained 24%, 31%, and 20% of the variance in analyses i, ii, and iii, respectively, and PC2 explained 19%, 26%, and 12%. PC1 scores were compared among genotypes by one-way ANOVA with Tukey’s post hoc test. No values were missing from the final inputs.

#### Persistence of Day 1 behavioral profiles

To test whether a mouse’s D1 profile persisted, the same 17 measures were calculated for each later session (D2, D3, D4, W2, W3, and W4). Each later-session dataset was z-scored across mice within that session and projected onto the D1 principal components; the D1 PCA was not refitted. Mice were matched across sessions by ID, and all 86 mice contributed to every session. PC1 scores across sessions were compared by two-way repeated-measures ANOVA (genotype × session; Fig. 2m); because each session was z-scored separately, this analysis tests genotype differences and genotype × session interactions rather than overall changes across sessions.

Individual stability of PC1 was assessed by simple linear regression of early (mean of D1 and W2) on late (mean of W3 and W4) scores (Fig. 2n).

To test persistence against chance, the Pearson correlation between each mouse’s D1 PC1 score and its PC1 score at each later session was compared with a null distribution (Fig. 2o). For each session, each measure was independently permuted across mice, which preserves each measure’s distribution but removes its covariance with the other measures of the same mouse; the permuted data were scaled and projected in the same way, and the correlation with D1 PC1 was recalculated. This was repeated 1,000 times per session, and the permutation p value was (1 + the number of permutations with a correlation at least as high as observed)/1,001, so the smallest attainable value was p ≈ 0.001.

### Genotype classification by assay

To determine which assays carried information about genotype (Fig. 5t), logistic-regression classifiers (scikit-learn; L2 penalty, C = 0.5) were trained to distinguish wild-type from homozygous mice using the measures from each assay separately (novel dyadic test, 8 measures; D1 sibling group, 17; three-chamber test, 2; open field, 2; elevated mazes, 2) and from all 31 measures combined; measures were z-scored within each training fold. Accuracy was estimated by leave-one-cage-out cross-validation, so that each mouse was classified by a model trained without its cagemates, and is reported as balanced accuracy (the mean of the wild-type and homozygous correct-classification rates; chance = 50%). For the combined model, significance was assessed against 200 permutations of the genotype labels.

## QUANTIFICATION AND STATISTICAL ANALYSIS

Analyses were performed in GraphPad Prism (v11.2), with additional processing in Python (v3.8.5; NumPy 1.24.4, pandas 2.0.3, SciPy 1.10.1, Shapely 1.8.4, NetworkX 2.5, scikit-learn 1.3.1) and R (v4.4.2). Genotype comparisons used one-way ANOVA with Tukey’s post hoc tests, except initiated chase time, chase bouts, and object contact in the novel dyadic test (Fig. 3l–n), which used Kruskal–Wallis tests with Dunn’s multiple comparisons. Measures with more than one level per mouse (contact or spatial categories, time bins, chambers, and Avg versus Chase speed) used two-way repeated-measures ANOVA with Tukey’s post hoc tests, or Fisher’s LSD for Fig. 3e, h, i and Fig. S1e. Longitudinal data used two-way repeated-measures ANOVA (genotype × session). Familiar-versus-novel comparisons used two-way repeated-measures ANOVA (genotype × partner) with Tukey’s post hoc tests, comparing genotypes within each partner condition (Fig. 4) and partners within each genotype (Fig. S7). Estrous effects were tested by one-way ANOVA across the four stages (genotypes pooled) and by two-way ANOVA (genotype × P/E versus DI/DII) with Tukey’s post hoc tests. Sex comparisons used two-way ANOVA (sex × genotype) with Tukey’s post hoc tests, or uncorrected Fisher’s LSD for Fig.

S6m–o and S8i–l. The proportions of mice that chased were compared by χ² test (Fig. 3j), and stability by simple linear regression and Pearson correlation. Persistence of D1 profiles was tested by permutation, and genotype classification by cross-validated logistic regression (see above). Statistics for Fig. 2k–o and 5q–t were computed in Python and checked against Prism. Data are presented as mean ± SEM unless otherwise stated.

Unless otherwise stated, n refers to individual mice; pairwise analyses use cagemate pairs as the unit. Exact n, the statistical test, and the definition of center and dispersion for each panel are given in the figure legends. All tests were two-tailed with α = 0.05; *p < 0.05, **p < 0.01, ***p < 0.001, ****p < 0.0001; #0.05 ≤ p < 0.10; where a trend is reported, the exact p value is given in the figure or legend. Data were excluded only as described above (insufficient estrous smears; mouse 1404, a wild-type female euthanized before completing all assays, from the three-chamber, open-field, and elevated-maze analyses (Fig. 4i–p, 5, S7i–q, S8, and S9); and one cage of four mice (1306–1309), excluded from all analyses because one cagemate required euthanasia, which altered the group’s social dynamics).

*Videos S1 and S2 are provided as separate files*.

---

**Figure S1.**
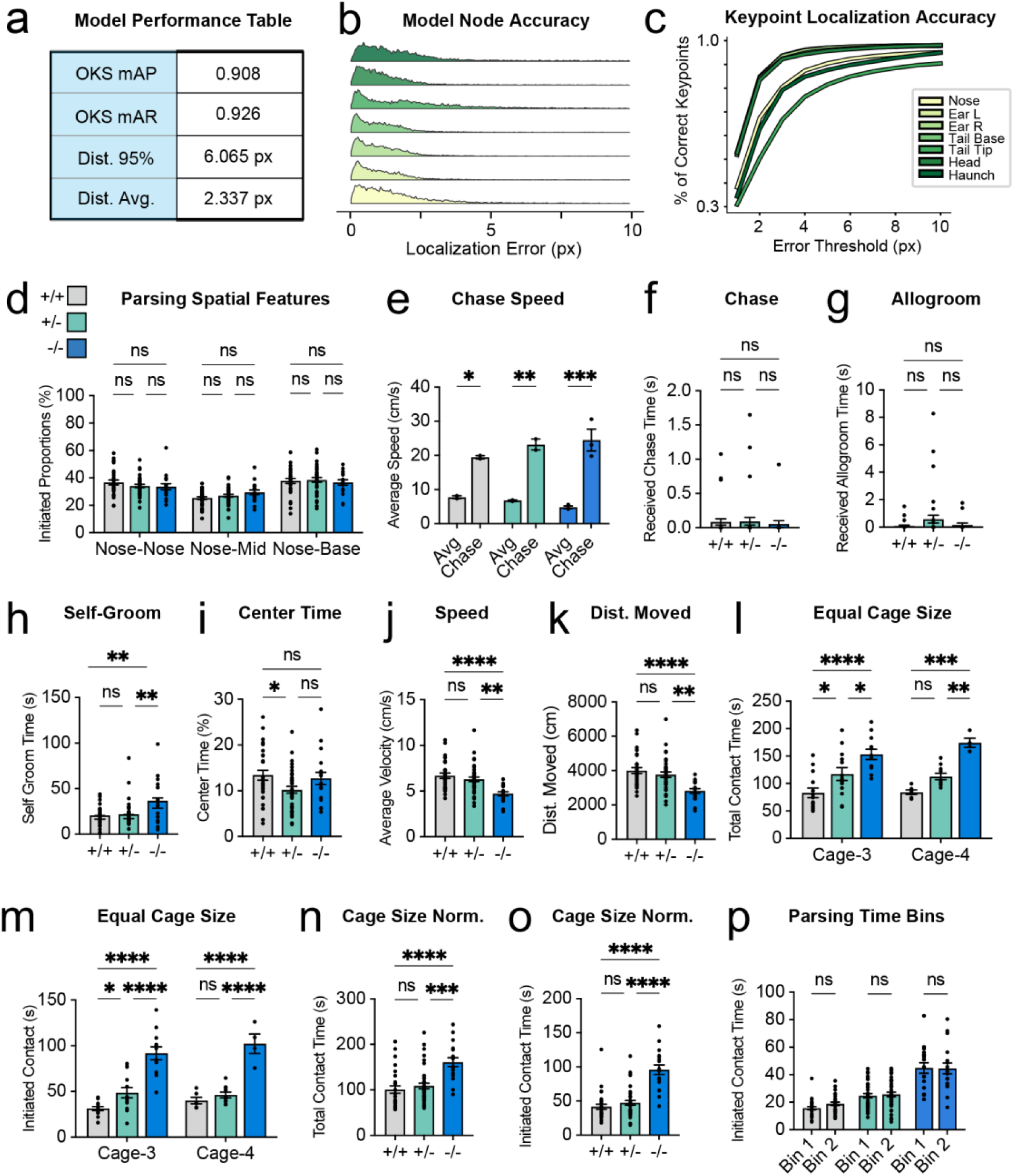
SLEAP model performance and single-session sibling-group measures, related to Figure 1. (A) Model performance on held-out validation frames: object keypoint similarity mean average precision (OKS mAP), 0.908; OKS mean average recall (mAR), 0.926; 95th-percentile localization error, 6.065 px; mean localization error, 2.337 px (1,200 × 1,200 px frames). (B) Distribution of localization error for each keypoint. (C) Percentage of correct keypoints as a function of error threshold for each keypoint. (D) Proportion of initiated contact by spatial type. (E) Average speed overall (Avg) and during chasing (Chase) in mice that initiated chasing (n = 2 WT, 2 HET, and 3 KO). (F) Received chase time. (G) Received allogrooming time. (H) Self-grooming time. (I) Percentage of time in the arena center. (J) Average speed. (K) Distance moved. (L and M) Total (L) and initiated (M) contact time (s) restricted to cages of three or four mice. (N and O) Total (N) and initiated (O) contact time normalized for cage size (contact time divided by the number of cagemates, cage size − 1). (P) Initiated contact time in the first (bin 1) and second (bin 2) 5 min of the session. Data are represented as mean ± SEM with individual mice as dots. n as in Figure 1. Two-way repeated-measures ANOVA with Tukey’s post hoc test (D); two-way repeated-measures ANOVA with Fisher’s LSD test (E); one-way ANOVA with Tukey’s post hoc test (F–K, N, O); two-way ANOVA (genotype × cage size) with Tukey’s post hoc test (L, M); two-way repeated-measures ANOVA with Tukey’s post hoc test, bin 1 versus bin 2 within genotype (P). Brackets: long bar, WT versus KO; short bars, WT versus HET (left) and HET versus KO (right). *p < 0.05, **p < 0.01, ***p < 0.001, ****p < 0.0001; ns, not significant.

**Figure S2.**
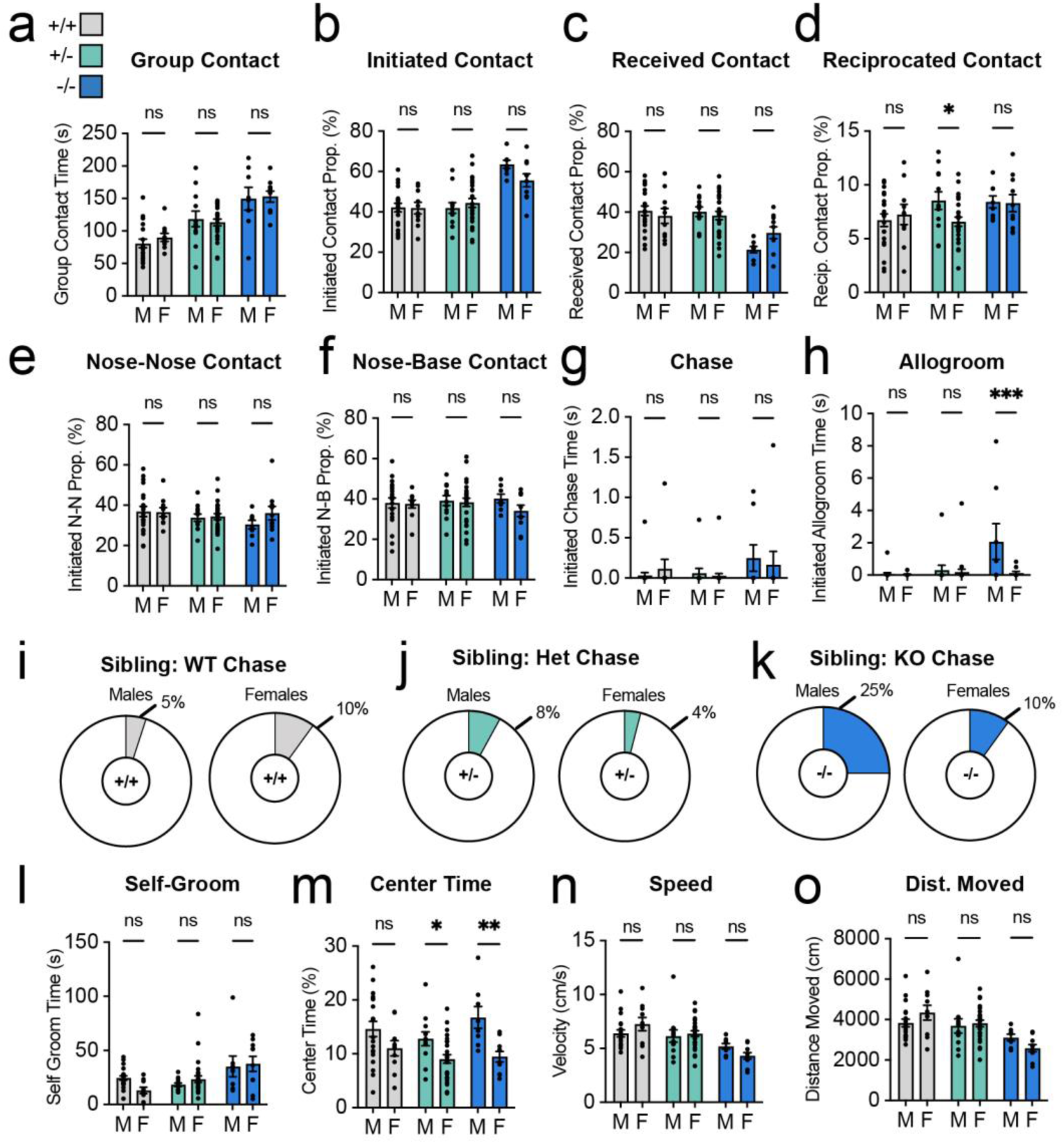
Sex comparisons for single-session sibling-group behavior, related to Figure 1. Males (M) and females (F) were compared within each genotype. (A) Group contact time. (B) Percentage of contact initiated. (C) Percentage of contact received. (D) Percentage of contact reciprocated. (E) Percentage of initiated contact that was nose-to-nose. (F) Percentage of initiated contact that was nose-to-base. (G) Initiated chase time. (H) Initiated allogrooming time. (I–K) Percentage of males and females that initiated chasing among WT (I; males 5%, 1 of 20; females 10%, 1 of 10), HET (J; males 8%, 1 of 12; females 4%, 1 of 26), and KO (K; males 25%, 2 of 8; females 10%, 1 of 10) mice. (L) Self-grooming time. (M) Percentage of time in the center. (N) Average speed. (O) Distance moved. Data are represented as mean ± SEM with individual mice as dots. n = 20 males and 10 females (WT), 12 males and 26 females (HET), and 8 males and 10 females (KO). Two-way ANOVA (sex × genotype) with Tukey’s post hoc test (A–H, L–O). *p < 0.05, **p < 0.01, ***p < 0.001, ****p < 0.0001; ns, not significant.

**Figure S3.**
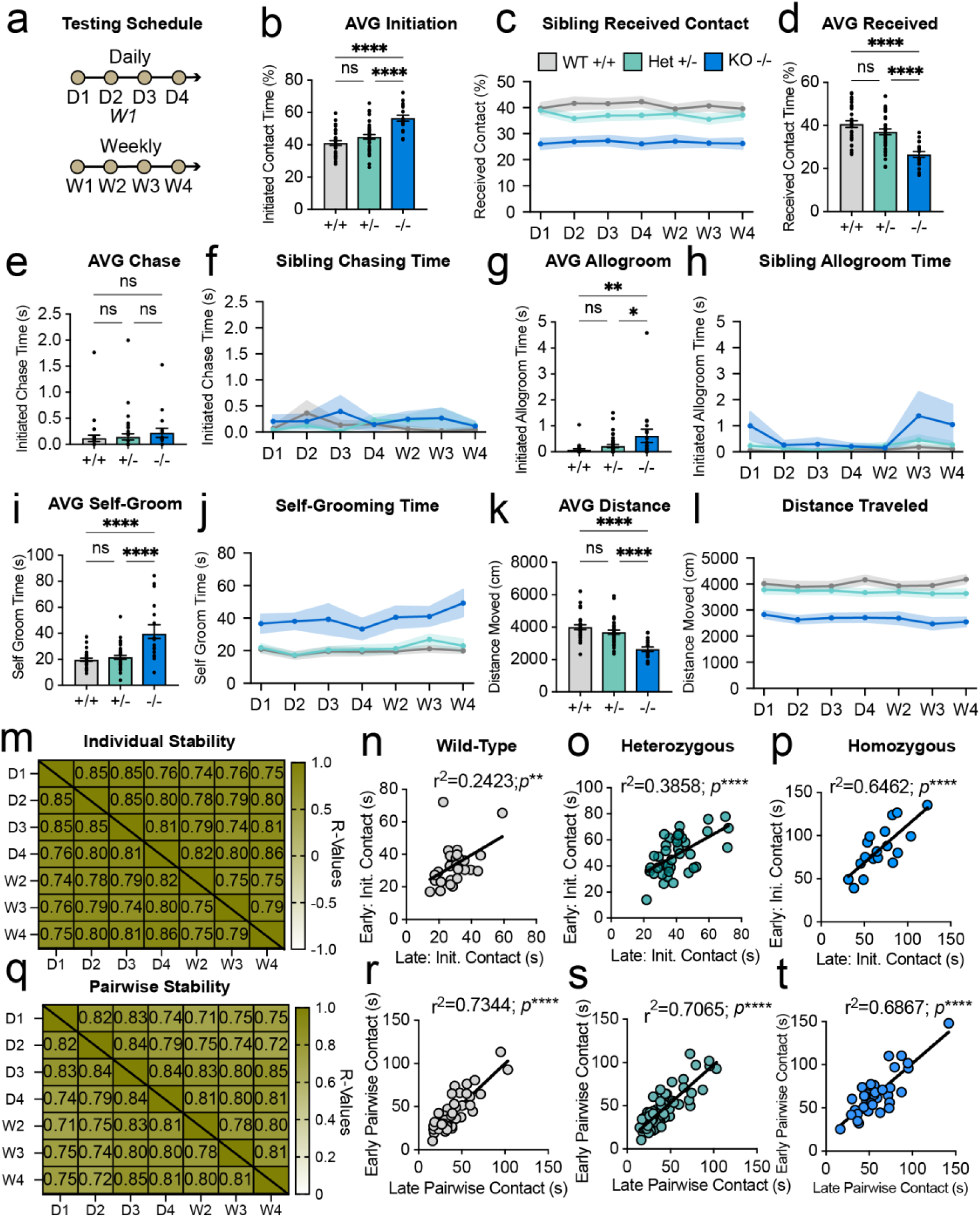
Additional measures from chronic sibling-group tracking, related to Figure 2. (A) Testing schedule. (B) Percentage of contact initiated, averaged across sessions. (C and D) Percentage of contact received across sessions (C) and averaged across sessions (D). (E and F) Initiated chase time averaged across sessions (E) and at each session (F). (G and H) Initiated allogrooming time averaged across sessions (G) and at each session (H). (I and J) Self-grooming time averaged across sessions (I) and at each session (J). (K and L) Distance moved averaged across sessions (K) and at each session (L). (M) Correlation (r) of individual initiated contact between every pair of sessions. (N–P) Early versus late initiated contact for WT (N; r² = 0.2423, p < 0.01), HET (O; r² = 0.3858, p < 0.0001), and KO (P; r² = 0.6462, p < 0.0001) mice. (Q) Correlation (r) of pairwise total contact between every pair of sessions. (R–T) Early versus late pairwise contact for WT (R; r² = 0.7344), HET (S; r² = 0.7065), and KO (T; r² = 0.6867) pairs; all p < 0.0001. Each cagemate pair is shown in the panel for each genotype it contains (51, 79, and 35 pairs in R, S, and T; 105 unique pairs). Data are represented as mean ± SEM with individual mice as dots; lines and shading show mean ± SEM; lines in (N)–(P) and (R)–(T) are simple linear regressions. n as in Figure 2. Early, mean of D1 and W2; late, mean of W3 and W4. One-way ANOVA with Tukey’s post hoc test (B, D, E, G, I, K); two-way repeated-measures ANOVA (genotype × session) (C, F, H, J, L); Pearson correlation (M, Q); simple linear regression (N–P, R–T). Brackets: long bar, WT versus KO; short bars, WT versus HET (left) and HET versus KO (right). *p < 0.05, **p < 0.01, ***p < 0.001, ****p < 0.0001; ns, not significant.

**Figure S4.**
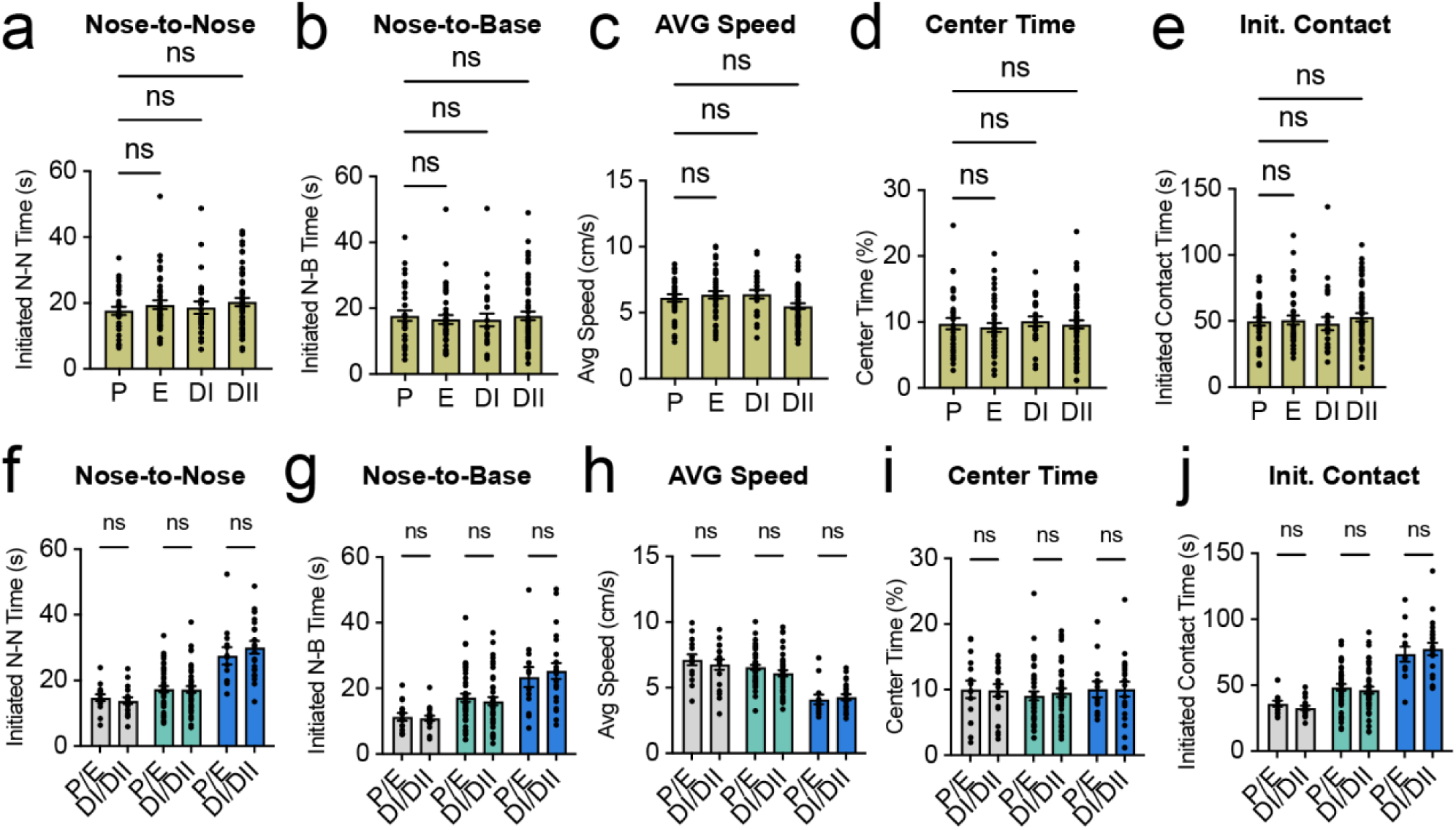
Estrous stage does not affect social or non-social measures in females, related to Figure 2. (A–E) Initiated nose-to-nose time (A), initiated nose-to-base time (B), average speed (C), percentage of time in the center (D), and initiated contact time (E) by estrous stage (P, proestrus; E, estrus; DI, diestrus I; DII, diestrus II), pooled across genotypes. (F–J) The same measures with observations grouped as P/E or DI/DII within each genotype. Data are represented as mean ± SEM with individual observations (sessions) as dots. n = 46 females; 33 P, 40 E, 26 DI, and 58 DII sessions; 157 total observations; 15 and 19 WT, 45 and 43 HET, and 13 and 22 KO observations in P/E and DI/DII, respectively (F–J). One-way ANOVA with Tukey’s post hoc test (A–E); two-way ANOVA with Tukey’s post hoc test, P/E versus DI/DII within genotype (F–J). ns, not significant.

**Figure S5.**
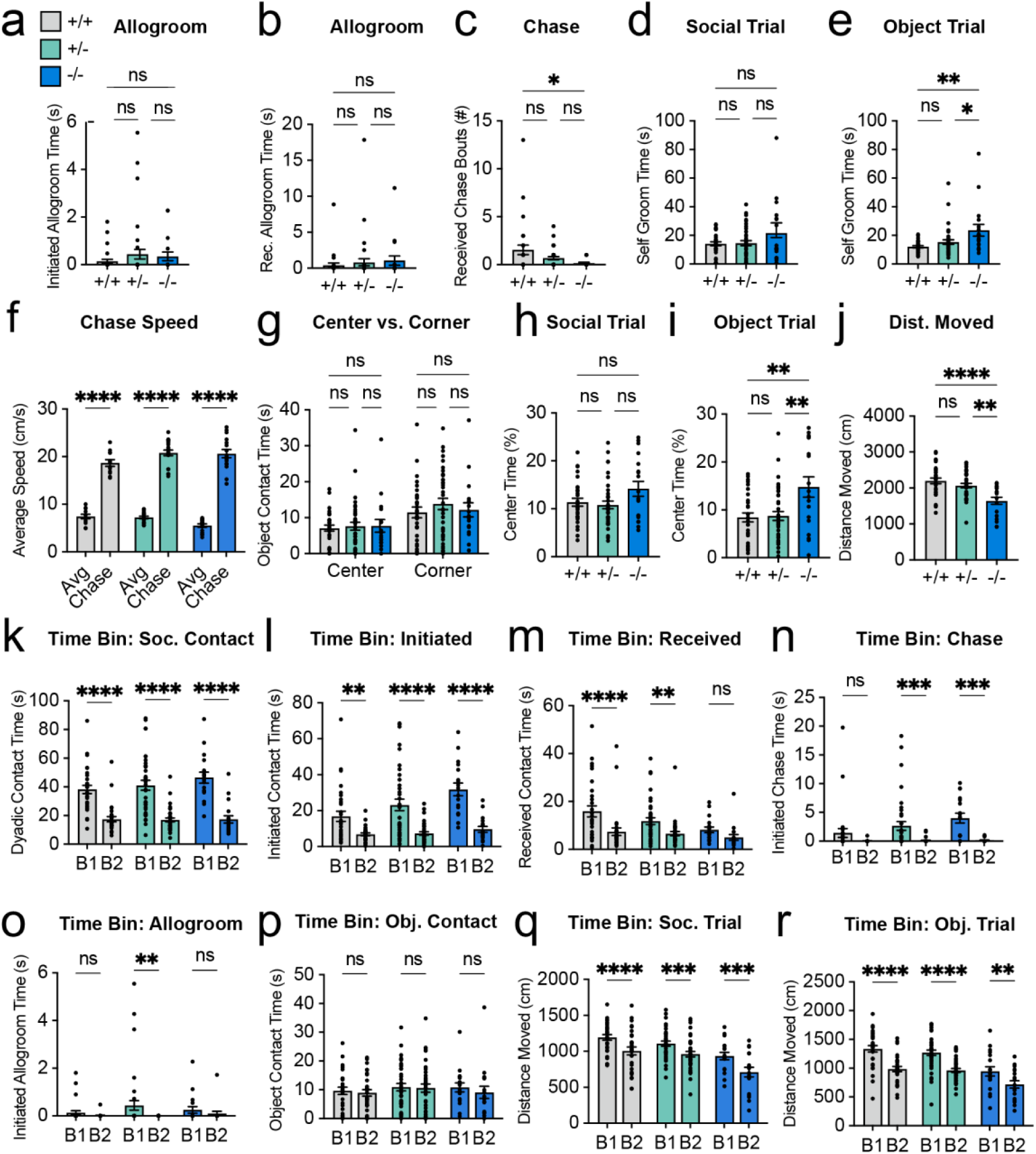
Additional measures and time-binned analysis of the novel dyadic test, related to Figure 3. (A) Initiated allogrooming time. (B) Received allogrooming time. (C) Number of chase bouts received from the stranger. (D and E) Self-grooming time during the social (D) and object (E) trials. (F) Average speed overall (Avg) and during chasing (Chase) in subjects that initiated chasing (n = 12 WT, 21 HET, and 16 KO). (G) Time contacting the center and corner objects. (H and I) Percentage of time in the center during the social (H) and object (I) trials. (J) Distance moved during the social trial. (K–R) Comparison of the first (B1) and second (B2) halves of each 5 min trial: dyadic contact time (K), initiated contact time (L), received contact time (M), initiated chase time (N), initiated allogrooming time (O), object contact time (P), and distance moved during the social (Q) and object (R) trials. Data are represented as mean ± SEM with individual mice as dots. n as in Figure 3. One-way ANOVA with Tukey’s post hoc test (A–E, H–J); two-way repeated-measures ANOVA with Tukey’s post hoc test (F, G, K–R). Brackets: long bar, WT versus KO; short bars, WT versus HET (left) and HET versus KO (right). *p < 0.05, **p < 0.01, ***p < 0.001, ****p < 0.0001; ns, not significant.

**Figure S6.**
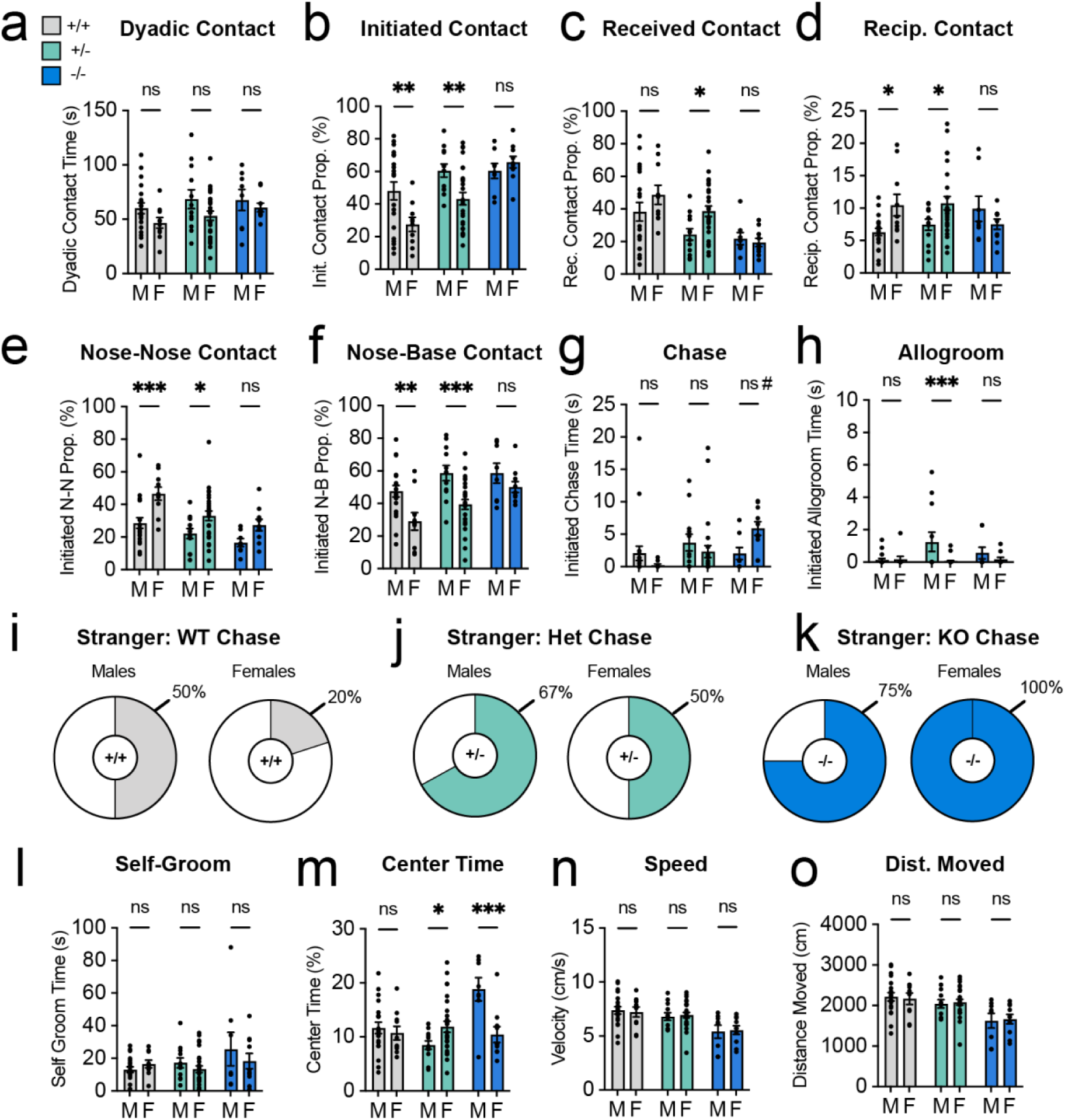
Sex comparisons for the novel dyadic test, related to Figure 3. Males (M) and females (F) were compared within each genotype. (A) Dyadic contact time. (B) Percentage of contact initiated. (C) Percentage of contact received. (D) Percentage of contact reciprocated. (E) Percentage of initiated contact that was nose-to-nose. (F) Percentage of initiated contact that was nose-to-base. (G) Initiated chase time. (H) Initiated allogrooming time. (I–K) Percentage of males and females that initiated chasing the stranger among WT (I; males 50%, 10 of 20; females 20%, 2 of 10), HET (J; males 67%, 8 of 12; females 50%, 13 of 26), and KO (K; males 75%, 6 of 8; females 100%, 10 of 10) subjects. (L) Self-grooming time. (M) Percentage of time in the center. (N) Average speed. (O) Distance moved. Data are represented as mean ± SEM with individual mice as dots. n = 20 males and 10 females (WT), 12 males and 26 females (HET), and 8 males and 10 females (KO). Two-way ANOVA (sex × genotype) with Tukey’s post hoc test (A–H, L); two-way ANOVA (sex × genotype) with uncorrected Fisher’s LSD test (M–O). #p < 0.10, *p < 0.05, **p < 0.01, ***p < 0.001, ****p < 0.0001; ns, not significant.

**Figure S7.**
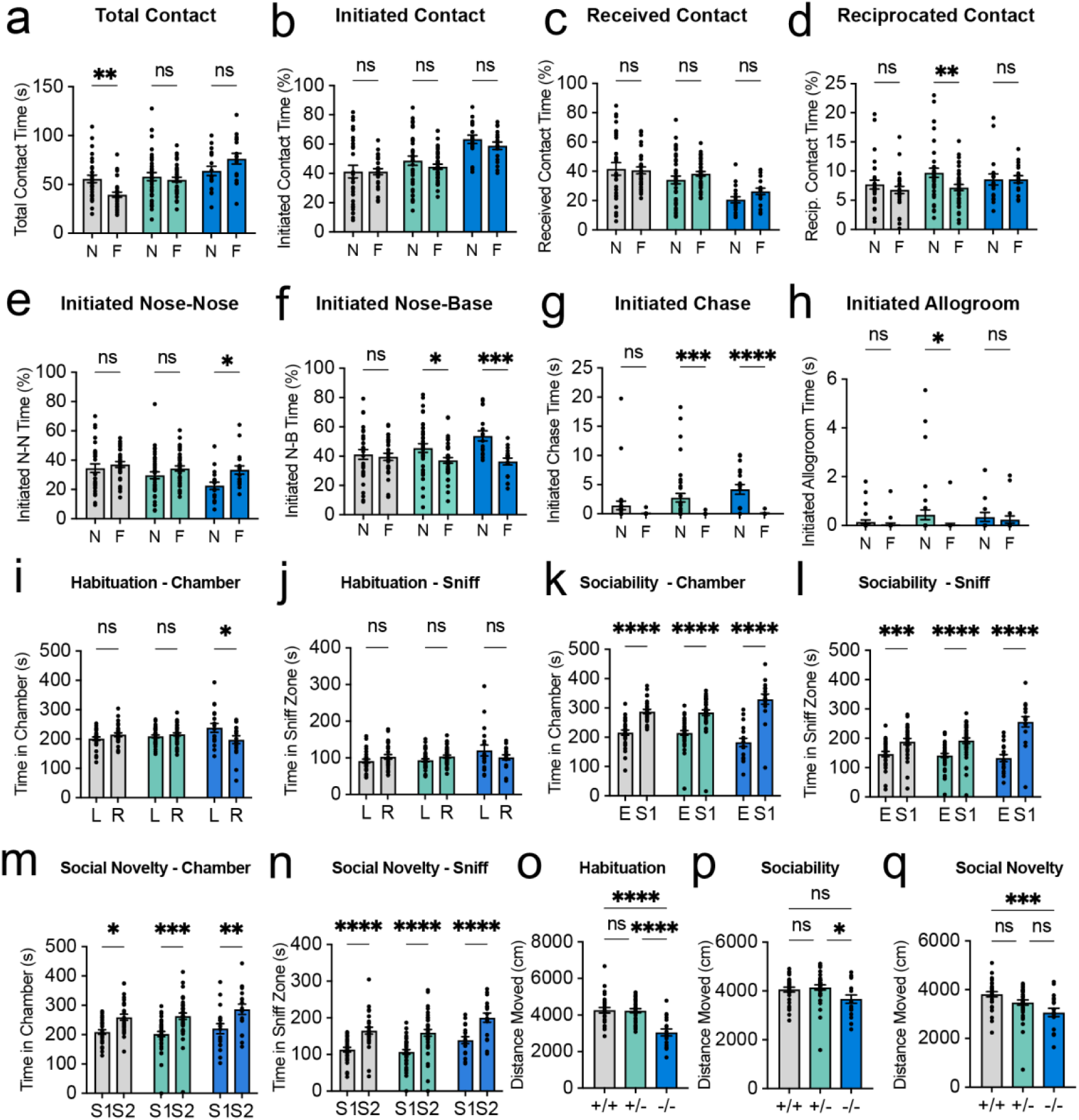
Novel-versus-familiar comparisons and three-chamber measures, related to Figure 4. (A–H) Novel (N) and familiar (F) partners compared within each genotype: total contact time (A), percentage of contact initiated (B), percentage of contact received (C), percentage of contact reciprocated (D), percentage of initiated contact that was nose-to-nose (E) or nose-to-base (F), initiated chase time (G), and initiated allogrooming time (H). (I and J) Three-chamber habituation: time in the left (L) and right (R) chambers (I) and sniff zones (J). (K and L) Sociability phase: time in the chamber (K) and sniff zone (L) of the empty enclosure (E) versus stranger 1 (S1). (M and N) Social novelty phase: time in the chamber (M) and sniff zone (N) of S1 versus stranger 2 (S2). (O–Q) Distance moved during habituation (O), sociability (P), and social novelty (Q). Data are represented as mean ± SEM with individual mice as dots. Contact times are raw values (no cage-size normalization). n as in Figure 4. Two-way repeated-measures ANOVA (genotype × partner) with Tukey’s post hoc test, novel versus familiar within genotype (A–H); two-way repeated-measures ANOVA with Tukey’s post hoc test, left versus right, E versus S1, or S1 versus S2 within genotype (I–N); one-way ANOVA with Tukey’s post hoc test (O–Q). Brackets: long bar, WT versus KO; short bars, WT versus HET (left) and HET versus KO (right). *p < 0.05, **p < 0.01, ***p < 0.001, ****p < 0.0001; ns, not significant.

**Figure S8.**
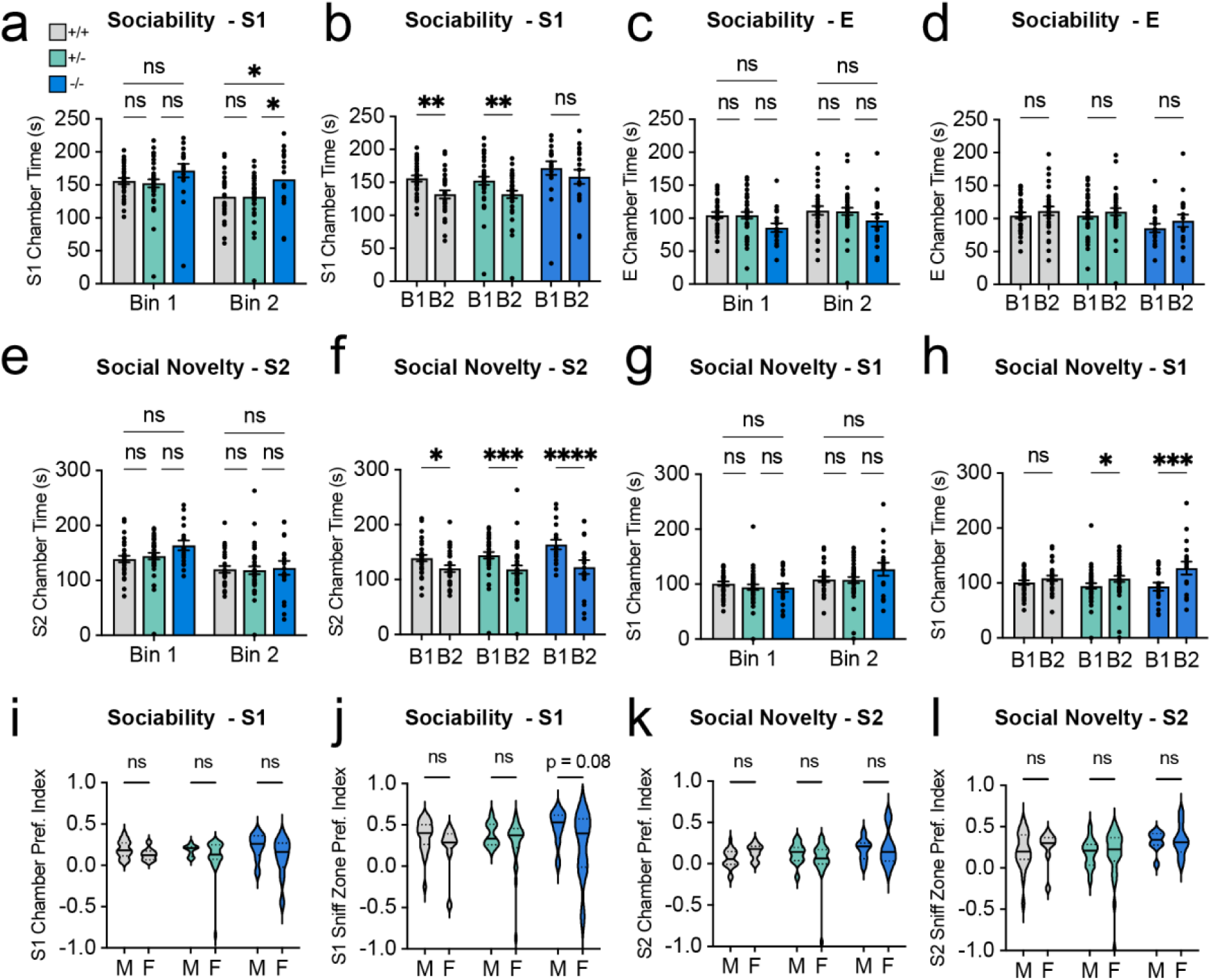
Time-binned three-chamber behavior and sex comparisons of preference indices, related to Figure 4. Each 10 min phase was divided into the first (bin 1, B1) and second (bin 2, B2) 5 min. (A and B) Sociability phase: time in the S1 chamber compared across genotypes within each bin (A) and between bins within each genotype (B). (C and D) Sociability phase: time in the empty-enclosure (E) chamber, by genotype within bin (C) and by bin within genotype (D). (E and F) Social novelty phase: time in the S2 chamber, by genotype within bin (E) and by bin within genotype (F). (G and H) Social novelty phase: time in the S1 chamber, by genotype within bin (G) and by bin within genotype (H). (I–L) Males (M) versus females (F) within each genotype: sociability preference index for chamber time (I) and sniff-zone time (J), and social novelty preference index for chamber time (K) and sniff-zone time (L). Data are represented as mean ± SEM with individual mice as dots; violin plots show the median (solid line) and quartiles (dotted lines). n as in Figure 4; for (I–L), 20 males and 9 females (WT), 12 males and 26 females (HET), and 8 males and 10 females (KO). Two-way repeated-measures ANOVA with Tukey’s post hoc test (A–H); two-way ANOVA (sex × genotype) with uncorrected Fisher’s LSD test (I–L); the exact p-value is shown for the KO comparison in (J). Brackets: long bar, WT versus KO; short bars, WT versus HET (left) and HET versus KO (right). *p < 0.05, **p < 0.01, ***p < 0.001, ****p < 0.0001; ns, not significant.

**Figure S9.**
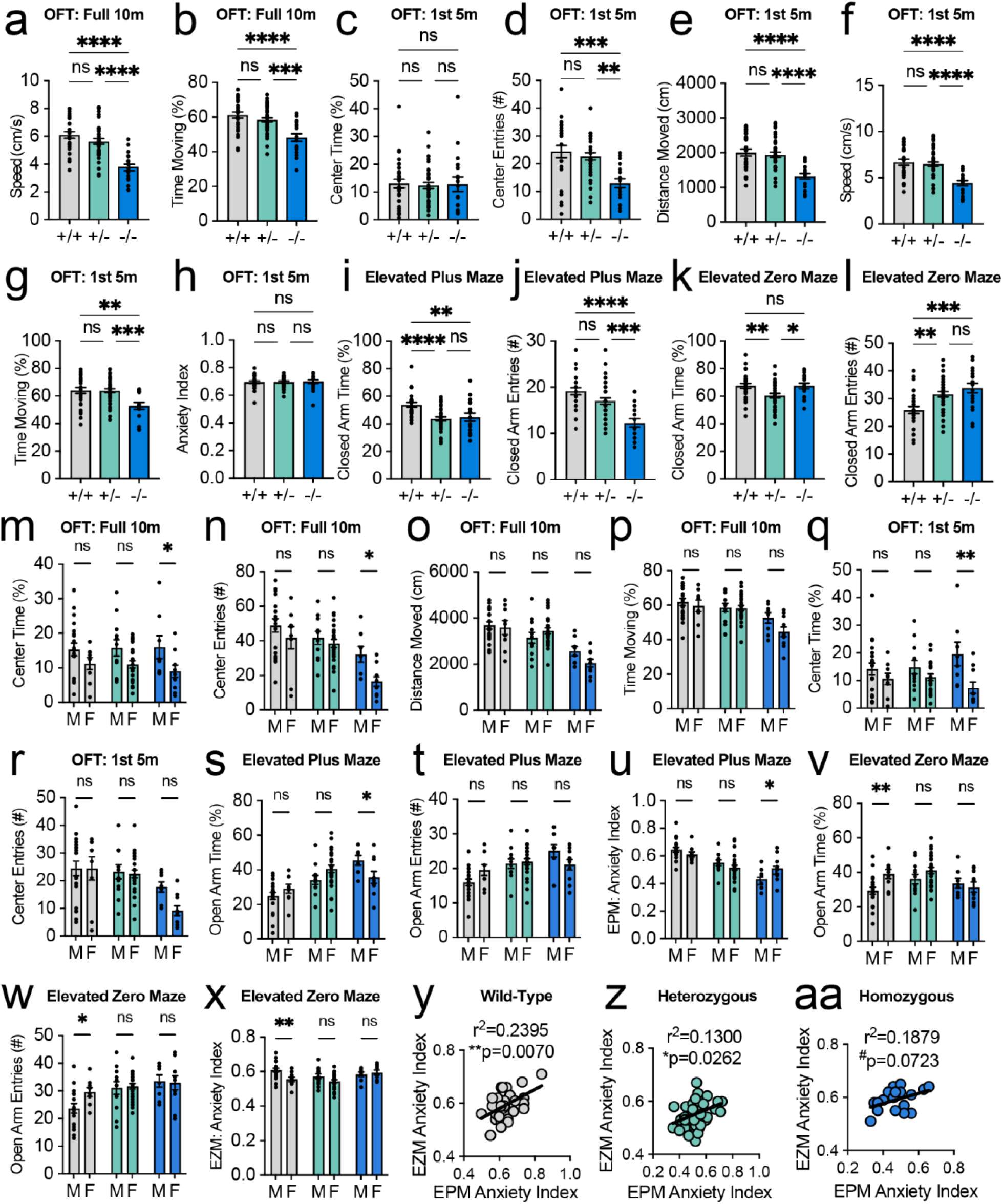
Additional anxiety-like and locomotor measures and sex comparisons, related to Figure 5. (A and B) OFT, full 10 min: speed (A) and percentage of time moving (B). (C–H) OFT, first 5 min: percentage of time in the center (C), center entries (D), distance moved (E), speed (F), percentage of time moving (G), and anxiety index (H; calculated as in Figure 5J from center time and center entries). (I and J) EPM: percentage of time in the closed arms (I) and closed-arm entries (J). (K and L) EZM: percentage of time in the closed quadrants (K) and closed-quadrant entries (L). (M–X) Males (M) versus females (F) within each genotype. OFT, full 10 min: center time (M), center entries (N), distance moved (O), and time moving (P). OFT, first 5 min: center time (Q) and center entries (R). EPM: open-arm time (S), open-arm entries (T), and anxiety index (U). EZM: open-quadrant time (V), open-quadrant entries (W), and anxiety index (X). (Y–AA) Correlation between EPM and EZM anxiety indices within WT (Y; r² = 0.2395, p = 0.0070), HET (Z; r² = 0.1300, p = 0.0262), and KO (AA; r² = 0.1879, p = 0.0723) mice. Data are represented as mean ± SEM with individual mice as dots; lines in (Y–AA) are linear regressions. n as in Figure 5; for (M–X), 20 males and 9 females (WT), 12 males and 26 females (HET), and 8 males and 10 females (KO). One-way ANOVA with Tukey’s post hoc test (A–L); two-way ANOVA (sex × genotype) with Tukey’s post hoc test (M–X); simple linear regression (Y–AA). Brackets: long bar, WT versus KO; short bars, WT versus HET (left) and HET versus KO (right). #p < 0.10, *p < 0.05, **p < 0.01, ***p < 0.001, ****p < 0.0001; ns, not significant.

**Supplementary Table S1.**
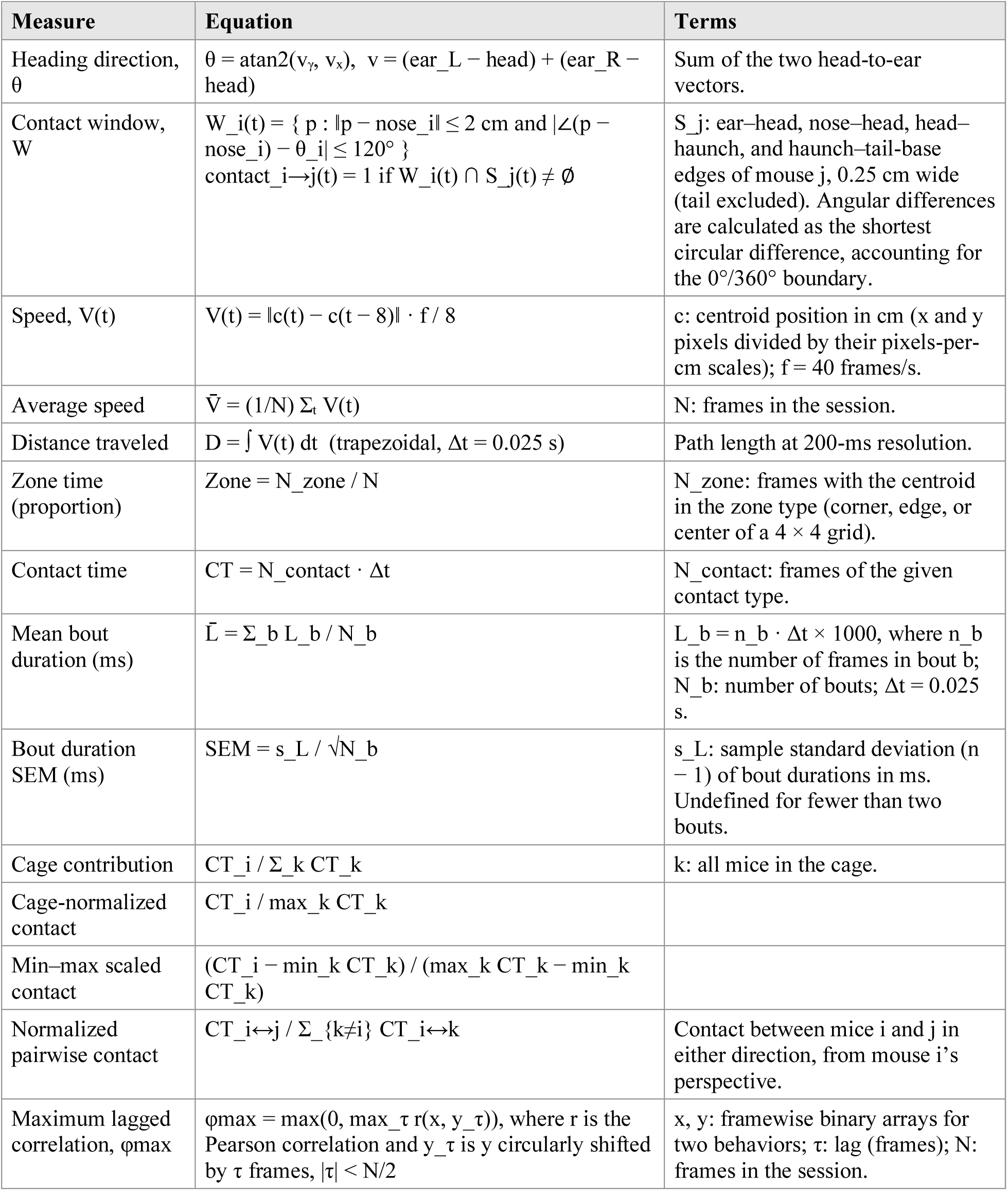
Equations for SLEAP-derived measures.

| Measure | Equation | Terms |
|---|---|---|
| Heading direction, θ | θ = atan2(v<sub>γ</sub>, v<sub>x</sub>), v = (ear_L − head) + (ear_R − head) | Sum of the two head-to-ear vectors. |
| Contact window, W | W_i(t) = { p : ‖p − nose_i‖ ≤ 2 cm and \|∠(p − nose_i) − θ_i\| ≤ 120° }<br>contact_i→j(t) = 1 if W_i(t) ∩ S_j(t) ≠ ∅ | S_j: ear–head, nose–head, head–haunch, and haunch–tail-base edges of mouse j, 0.25 cm wide (tail excluded). Angular differences are calculated as the shortest circular difference, accounting for the 0°/360° boundary. |
| Speed, V(t) | V(t) = ‖c(t) − c(t − 8)‖ · f / 8 | c: centroid position in cm (x and y pixels divided by their pixels-per-cm scales); f = 40 frames/s. |
| Average speed | V̄ = (1/N) Σ<sub>t</sub> V(t) | N: frames in the session. |
| Distance traveled | D = ∫ V(t) dt (trapezoidal, Δt = 0.025 s) | Path length at 200-ms resolution. |
| Zone time (proportion) | Zone = N_zone / N | N_zone: frames with the centroid in the zone type (corner, edge, or center of a 4 × 4 grid). |
| Contact time | CT = N_contact · Δt | N_contact: frames of the given contact type. |
| Mean bout duration (ms) | L̄ = Σ_b L_b / N_b | L_b = n_b · Δt × 1000, where n_b is the number of frames in bout b; N_b: number of bouts; Δt = 0.025 s. |
| Bout duration SEM (ms) | SEM = s_L / √N_b | s_L: sample standard deviation (n − 1) of bout durations in ms. Undefined for fewer than two bouts. |
| Cage contribution | CT_i / Σ_k CT_k | k: all mice in the cage. |
| Cage-normalized contact | CT_i / max_k CT_k |  |
| Min–max scaled contact | (CT_i − min_k CT_k) / (max_k CT_k − min_k CT_k) |  |
| Normalized pairwise contact | CT_i↔j / Σ_{k≠i} CT_i↔k | Contact between mice i and j in either direction, from mouse i's perspective. |
| Maximum lagged correlation, φmax | φmax = max(0, max_τ r(x, y_τ)), where r is the Pearson correlation and y_τ is y circularly shifted by τ frames, \|τ\| < N/2 | x, y: framewise binary arrays for two behaviors; τ: lag (frames); N: frames in the session. |

**Supplementary Table S2.** Measures entered in each principal-component analysis and genotype-decoding model. Measures were chosen by rules that did not use genotype (see note). Each measure was z-scored across mice before PCA; measures were not weighted. Variance explained by PC1 and PC2: Fig. 2k, 24% and 19%; Fig. 5q, 31% and 26%; Fig. 5s, 20% and 12%. n = 86 mice (Fig. 2k–o) and 85 mice (Fig. 5q–s; mouse 1404, euthanized before completing all assays, excluded); Fig. 5t, 29 WT and 18 KO mice.

| **Analysis** | **#** | **Measure** | **Spreadsheet column or formula** |
|---|---|---|---|
| **Fig. 2k–o: D1 sibling group (17 measures); later sessions (D2–W4) were scored on the same measures and projected onto the D1 PCA** |  |  |  |
|  | # | Metric | Column in the spreadsheet (or formula) |
| Amount of contact | 1 | Total contact (s) | Total Contact Time (s) |
| Amount of contact | 2 | Initiated contact (s) | Initiated Total Contact Time (s) |
| Amount of contact | 3 | Received contact (s) | Received Total Contact Time (s) |
| Role | 4 | Initiated (% of contact) | Initiated Total Contact Time (s) / Total Contact Time (s) × 100 |
| Role | 5 | Reciprocated (% of contact) | Reciprocated Total Contact Time (s) / Total Contact Time (s) × 100 |
| Location | 6 | Nose-mid (% of contact) | Total Nose-mid Contact Time (s) / Total Contact Time (s) × 100 |
| Location | 7 | Initiated nose-nose (% of initiated) | Initiated Nose-nose Contact Time (s) / Initiated Total Contact Time (s) × 100 |
| Location | 8 | Initiated nose-mid (% of initiated) | Initiated Nose-mid Contact Time (s) / Initiated Total Contact Time (s) × 100 |
| Location | 9 | Received nose-nose (% of received) | Received Nose-nose Contact Time (s) / Received Total Contact Time (s) × 100 |
| Location | 10 | Received nose-mid (% of received) | Received Nose-mid Contact Time (s) / Received Total Contact Time (s) × 100 |
| Bout length | 11 | Initiated nose-nose bout length (ms) | Initiated Nose-nose Avg Bout Dur (ms) |
| Bout length | 12 | Initiated nose-mid bout length (ms) | Initiated Nose-mid Avg Bout Dur (ms) |
| Bout length | 13 | Received nose-nose bout length (ms) | Received Nose-nose Avg Bout Dur (ms) |
| Bout length | 14 | Received nose-mid bout length (ms) | Received Nose-mid Avg Bout Dur (ms) |
| Bout length | 15 | Dual bout length (ms) | Dual Total Avg Bout Dur (ms) |
| Non-social | 16 | Corner time (%) | Time in Corner (%) |
| Non-social | 17 | Edge time (%) | Time at Edge (%) |
| **Fig. 5q,r: anxiety-like and locomotor (10 measures)** |  |  |  |
|  | # | Metric | Column in the spreadsheet |
| Solo OFT | 1 | Center frequency | Solo OFT Center Freq |
| Solo OFT | 2 | Center time (%) | Solo OFT Center Time (%) |
| Solo OFT | 3 | Distance moved (cm) | Solo OFT Dist Moved (cm) |
| Solo OFT | 4 | Corner time (%) | Solo OFT Corner Time (%) |
| EZM | 5 | Open quadrant frequency | EZM Open Arm Freq |
| EZM | 6 | Open quadrant time (%) | EZM Open Arm Time (%) |
| EPM | 7 | Open arm frequency | EPM Open Arm Freq |
| EPM | 8 | Open arm time (%) | EPM Open Arm Time (%) |
| EPM | 9 | Closed arm frequency | EPM Closed Arm Freq |
| EPM | 10 | Center time (%) | EPM Center Time (%) |
| **Fig. 5s,t: all assays (31 measures, unweighted); Fig. 5t decoding used each assay’s measures alone and all 31 combined** |  |  |  |
| Assay | # | Metric | Column in the spreadsheet |
| SNT | 1 | Total contact (s) | Total Contact Time (s) |
| SNT | 2 | Initiated contact (s) | Initiated Total Contact Time (s) |
| SNT | 3 | Initiated (% of contact) | SNT Initiated Total Contact Prop (%) |
| SNT | 4 | Initiated nose-nose (% of initiated) | SNT Init Nose-nose Contact Prop (%) |
| SNT | 5 | Initiated nose-base (% of initiated) | SNT Init Nose-base Contact Prop (%) |
| SNT | 6 | Initiated bout length (ms) | Initiated Total Avg Bout Dur (ms) |
| SNT | 7 | Initiated chase time (s) | Initiated Chase Time (s) |
| SNT | 8 | Speed (cm/s) | Average Speed (cm/s) |
| D1 group | 9 | 17 D1 measures listed above | As above |
| 3-chamber | 10 | Sniff zone preference index, S1 vs habituation baseline | 3-Chamber S1vE S1 Sniff PI from Baseline |
| 3-chamber | 11 | Sniff zone preference index, S2 vs habituation baseline | 3-Chamber S1vS2 S2 Sniff PI from Baseline |
| Solo OFT | 12 | Center time (%) | Solo OFT Center Time (%) |
| Solo OFT | 13 | Distance moved (cm) | Solo OFT Dist Moved (cm) |
| EPM | 14 | Open arm time (%) | EPM Open Arm Time (%) |
| EZM | 15 | Open quadrant time (%) | EZM Open Arm Time (%) |

Selection rules for SLEAP group measures: measures calculated from cagemates’ values (cage contribution, cage-normalized and min–max scaled contact), time in a group of k, bout SEMs, and measures that were mostly zero were excluded; measures that differed among cages of 2–5 mice (genotype-adjusted ANOVA, p < 0.05) were excluded, including dual, nose-to-nose, and nose-to-base contact as a percentage of total contact, speed, center time, and bout counts; and one of each near-duplicate pair (|r| > 0.9) was removed (e.g., received % of contact, r = 0.96 with initiated %). Anxiety measures: border measures, velocity, time moving, corner entries, closed-arm/quadrant measures other than EPM closed-arm frequency, and anxiety indices were removed as near-duplicates (|r| ≥ 0.92). Decoding assay groups in Fig. 5t: Stranger, SNT measures 1–8; Sibling, the 17 D1 measures; 3-Cham, three-chamber measures 10–11; OFT, measures 12–13; EZM/EPM, measures 14–15; All, all 31. Percentages computed from two columns are given as formulas. SNT, stranger (novel dyadic) test; NN, nose-to-nose; NM, nose-to-mid.

